# Multiple introductions, trade-associated connectivity, and mito-nuclear discordance reveal complex invasion dynamics of *Aedes albopictus* in Colombia

**DOI:** 10.64898/2026.06.15.732288

**Authors:** Juan S. Mantilla-Granados, Eliana Calvo-Tapiero, Karol Montilla-López, Myriam Lucia Velandia-Romero, Carlos Morales, Jorge De Las Salas-Ali, Cristian Javier Salcedo-Amortegui, Luz Stella Buitrago, Liliana Quintero, Guillermo Rúa, Jaime E. Castellanos

## Abstract

**Background:** *Aedes albopictus* is among the world’s most invasive mosquito species and an important vector of dengue, Zika, and chikungunya viruses. Its global spread has been strongly associated with human-mediated transport and international trade, particularly through commodities such as used tires and ornamental plants. However, integrative studies combining population genetics, microbial symbiosis, and trade connectivity remain limited in Latin America, constraining understanding of invasion dynamics and dispersal processes.

**Methods:** *Aedes albopictus* populations from five Colombian departments sampled between 2019 and 2023 were analyzed using eight microsatellite loci and a ∼1.3-kb mitochondrial COI fragment. Wolbachia infection and lineage composition (wAlbA/wAlbB) were evaluated by PCR, and arbovirus detection (DENV, CHIKV, ZIKV) was performed using multiplex RT-PCR. Nuclear and mitochondrial differentiation (FST, ΦST), mito-nuclear discordance, and trade-related connectivity were evaluated in relation to geographic, national transport, and international trade variables derived from customs databases.

**Results:** Microsatellite analyses revealed admixed but structured populations consistent with multiple introductions and contemporary gene flow. Colombian populations showed nuclear genetic affinities with Asian, European, and North American populations, supporting a complex invasion history involving multiple geographically distributed lineages. In contrast, mitochondrial COI data showed evidence of regional genetic structure and heterogeneous mito-nuclear discordance among several population pairs. Notably, nuclear and mitochondrial markers captured contrasting dimensions of the invasion process: nuclear differentiation was positively associated with international trade intensity, particularly shipment frequency and used tire importation, whereas mitochondrial differentiation retained stronger regional structure and showed no detectable association with trade-related variables. Wolbachia prevalence ranged from 34% to 100% across departments and showed exploratory patterns consistent with localized mitochondrial differentiation. Natural detection of DENV and CHIKV RNA in larvae provided evidence of local arbovirus circulation.

**Conclusions:** Integrating population genetics, trade connectivity, and Wolbachia screening supports a scenario in which the Colombian invasion of *Ae. albopictus* has been shaped by multiple introductions, contemporary human-mediated connectivity, and partially discordant mito-nuclear histories. These findings highlight how different genomic compartments retain complementary signatures of invasion dynamics, with contemporary trade-associated connectivity primarily reflected in nuclear structure and regional lineage persistence retained in mitochondrial variation.

**Author Summary:** The Asian tiger mosquito, *Aedes albopictus*, is one of the world’s most invasive mosquito vectors and continues to expand across Latin America through human transportation and trade networks. However, the processes shaping its spread in the region remain poorly understood. We combined population genetics, international trade data, Wolbachia screening, and arbovirus surveillance to investigate the invasion dynamics of *Ae. albopictus* in Colombia. Our results revealed evidence of multiple introductions and ongoing genetic admixture, with international trade connectivity emerging as an important predictor of contemporary nuclear genetic structure. In contrast, mitochondrial DNA retained stronger regional patterns, generating heterogeneous mito-nuclear discordance among populations. These findings suggest that different genomic compartments retain distinct signatures of the invasion process, with trade-associated connectivity reflected primarily in nuclear variation and stronger regional structure preserved in mitochondrial lineages. More broadly, our study highlights the complex invasion dynamics of *Ae. albopictus* in Latin America illustrates how integrating genetics and human connectivity data can improve understanding of invasive vector spread.

## INTRODUCTION

Aedes albopictus is one of the world’s most invasive mosquito species, having colonized all continents except Antarctica and becoming one of the most widely distributed arthropod vectors [1,2]. Although Aedes aegypti remains the principal vector of dengue (DENV), Zika (ZIKV), chikungunya (CHIKV), and urban yellow fever (YFV), the epidemiological importance of *Ae. albopictus* has increased due to its broad ecological tolerance and expanding geographic range [1,3–5]. Its ability to establish in urban, peri-urban, and rural environments has facilitated the emergence of autochthonous arboviral outbreaks in recently invaded regions, including Europe [5–8]. In addition, adaptive viral mutations such as CHIKV E1-A226V have further increased the relevance of some Ae. albopictus populations by enhancing transmission efficiency [7].

The rapid global expansion of *Ae. albopictus* has been accelerated by globalization, climate change, and increasing human mobility, with eggs and larvae readily dispersed through maritime, aerial, and terrestrial transportation networks, particularly via the international trade of used tires and ornamental plants [1,9]. Reconstructing invasion pathways and distinguishing between historical and contemporary dispersal processes therefore requires molecular markers capable of capturing different dimensions of population connectivity. Mitochondrial markers such as cytochrome oxidase I (COI) provide valuable information on haplotype diversity and broad-scale colonization history, particularly when long COI fragments are analyzed [10]. In contrast, nuclear microsatellites allow finer-scale inference of recent introductions, admixture, and ongoing gene flow [11,12]. The integration of mitochondrial and nuclear markers has therefore become increasingly important for reconstructing complex invasion dynamics and identifying how different spatial and temporal processes contribute to the spread of invasive mosquito populations.

Interactions between *Ae. albopictus* and microbial symbionts may also influence invasion dynamics and population genetic structure. *Wolbachia*, a maternally inherited bacterial endosymbiont naturally present in *Ae. albopictus* as the wAlbA and wAlbB lineages, is known to induce cytoplasmic incompatibility and can influence arbovirus susceptibility and reproductive dynamics [13,14]. Because both *Wolbachia* and mitochondrial DNA are maternally inherited, shifts in endosymbiont prevalence may alter mitochondrial diversity through cytoplasmic incompatibility, potentially obscuring demographic inferences based solely on mitochondrial markers [15]. Previous studies in Europe have further shown that multiple introductions, human-mediated dispersal, and heterogeneous *Wolbachia* dynamics can jointly structure *Ae. albopictus* populations, emphasizing the importance of integrative approaches combining population genetics, microbial symbiosis, and invasion ecology [16].

Despite the growing epidemiological importance of *Ae. albopictus*, integrative studies addressing invasion dynamics in Latin America remain limited. Most available studies have analyzed mitochondrial and nuclear markers independently, revealing evidence of regional connectivity among North, Central, and South American populations [7,10,17,18], but few have incorporated multilocus genetic data together with trade-related connectivity and endosymbiont dynamics. In Colombia, *Ae. albopictus* was first detected in the Amazon region in 1998 [19]and has since expanded across ecologically diverse regions of the country [5]. Reports of natural arboviral infections [20,21] and evidence of viral dissemination in Colombian populations [5] further underscore the epidemiological relevance of understanding population structure and invasion pathways in this emerging vector. However, population genetic data from Colombia remain scarce, limiting the ability to reconstruct introduction histories, evaluate lineage persistence, and understand how contemporary human-mediated connectivity shapes invasion dynamics.

Colombia’s geographic complexity, environmental heterogeneity, and intense connectivity through ports, airports, and terrestrial transportation corridors make it an ideal system for investigating mosquito invasion dynamics. Here, we integrated nuclear and mitochondrial population genetics, international trade-related connectivity data, and Wolbachia screening to investigate the invasion dynamics of *Ae. albopictus* and identify factors associated with its expansion across Colombia. We tested three main hypotheses: (1) Colombian populations reflect multiple introductions and complex admixture from genetically distinct sources; (2) population genetic differentiation is associated with major trade and transportation networks linked to human mobility; and (3) heterogeneous Wolbachia prevalence and lineage composition may contribute to localized mito–nuclear discordance. By integrating population genetics within ecological and anthropogenic contexts, this study provides a framework for understanding how trade-associated connectivity, microbial symbiosis, and demographic processes may contribute to shaping the spread and genetic structure of invasive vector populations.

## METHODS

### Study area and sampling design

This study was conducted in urban settings across 18 municipalities distributed among the departments of Antioquia, Cauca, Meta, Quindío, and Vichada, Colombia (Figure 1, Table 1). Sampling sites were selected based on both historical and recent records confirming the presence of *Ae. albopictus*. Larval collections were carried out as part of routine *Aedes* spp. surveillance programs implemented by local health authorities between 2019 and 2023.

**Figure 1.**
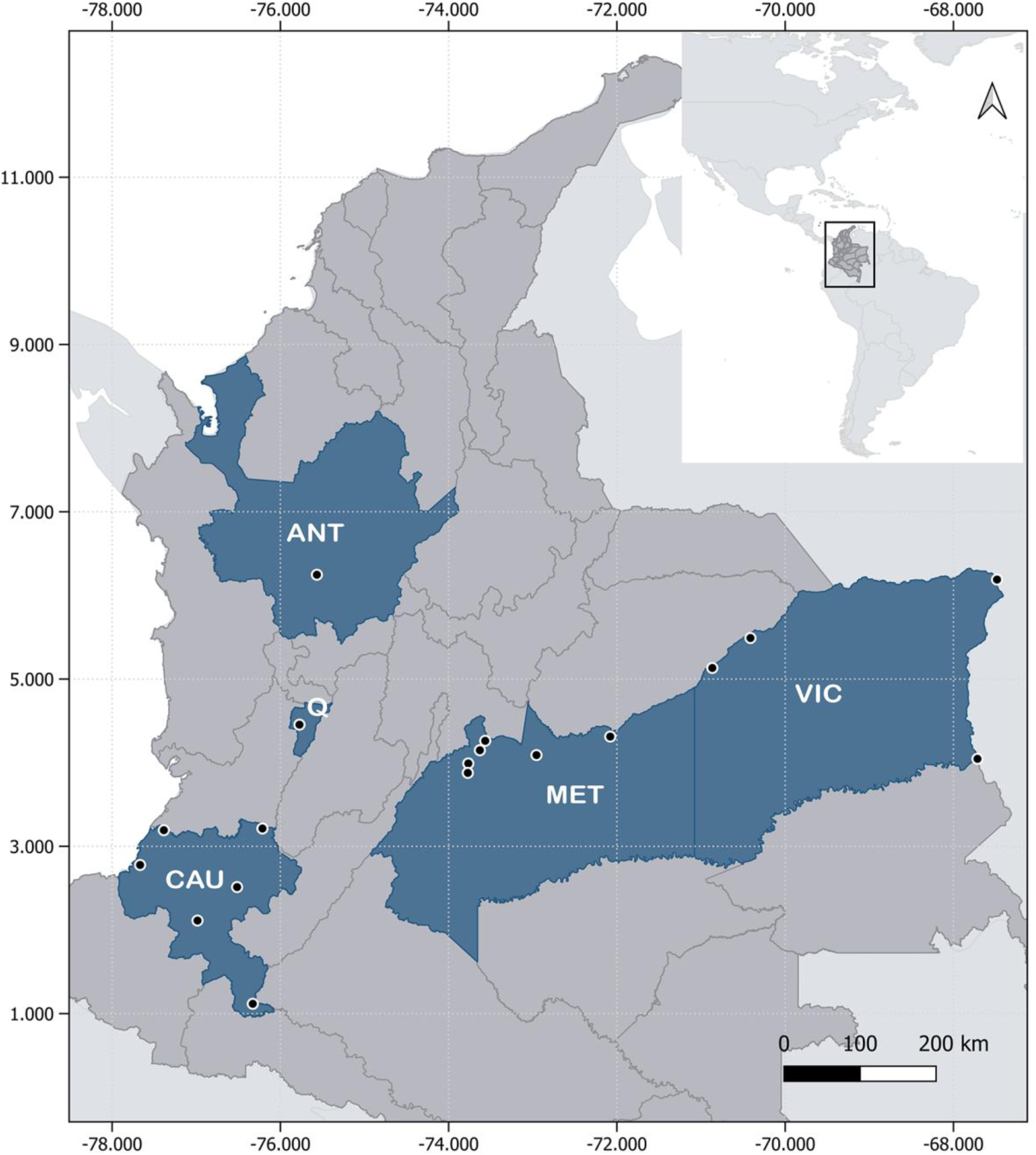
Map showing the location of the 18 sampled municipalities in Colombia. The Colombia departments are marked in gray and the sampled departments in blue Antioquia (ANT); Quindio (Q); Cauca (Cau), Meta (MET); Vichada (VIC). Black dots indicate the location of the sampled areas. Basemap layers for administrative division were obtained from the open data portal https://geoportal.dane.gov.co/.

**Table 1.**
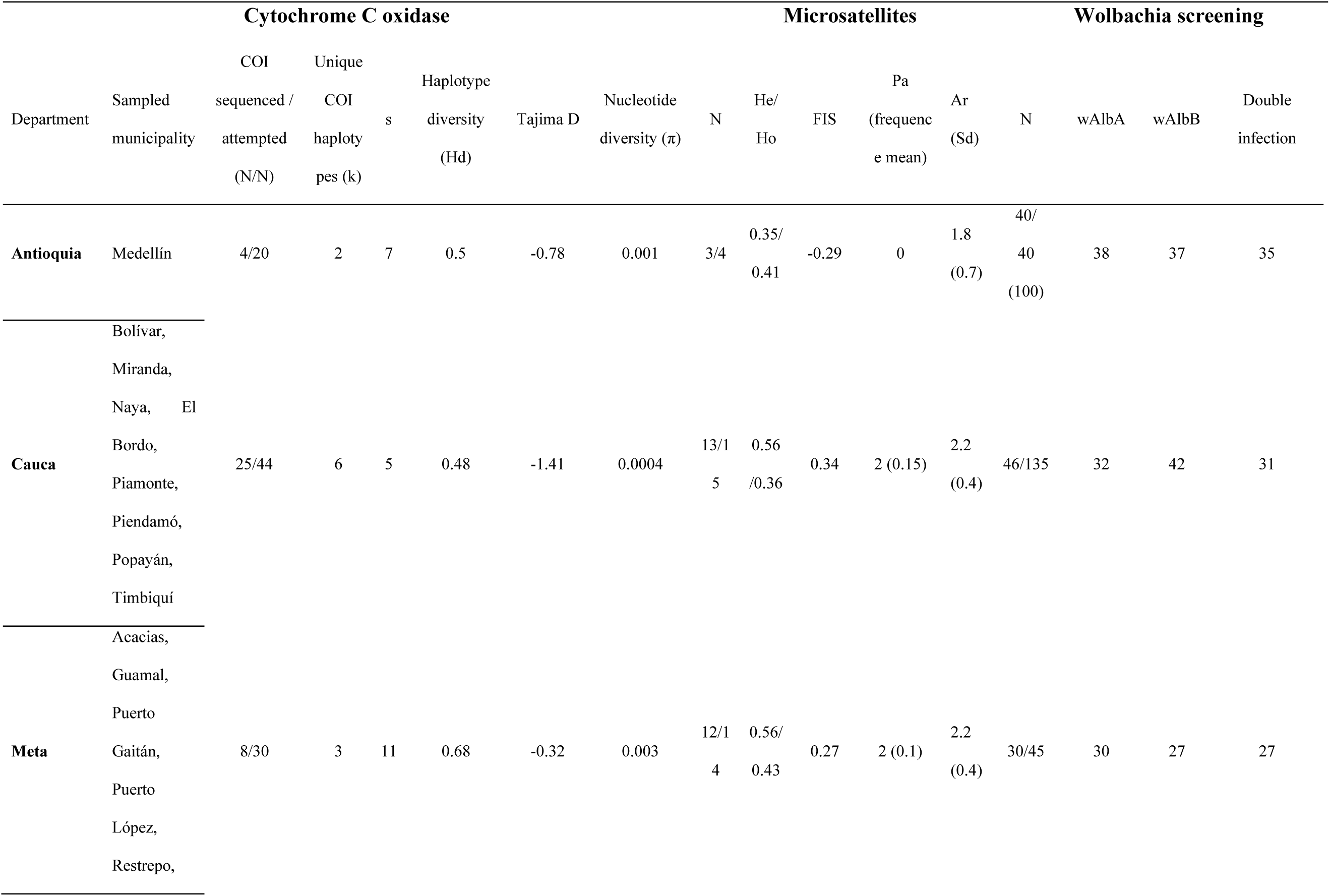
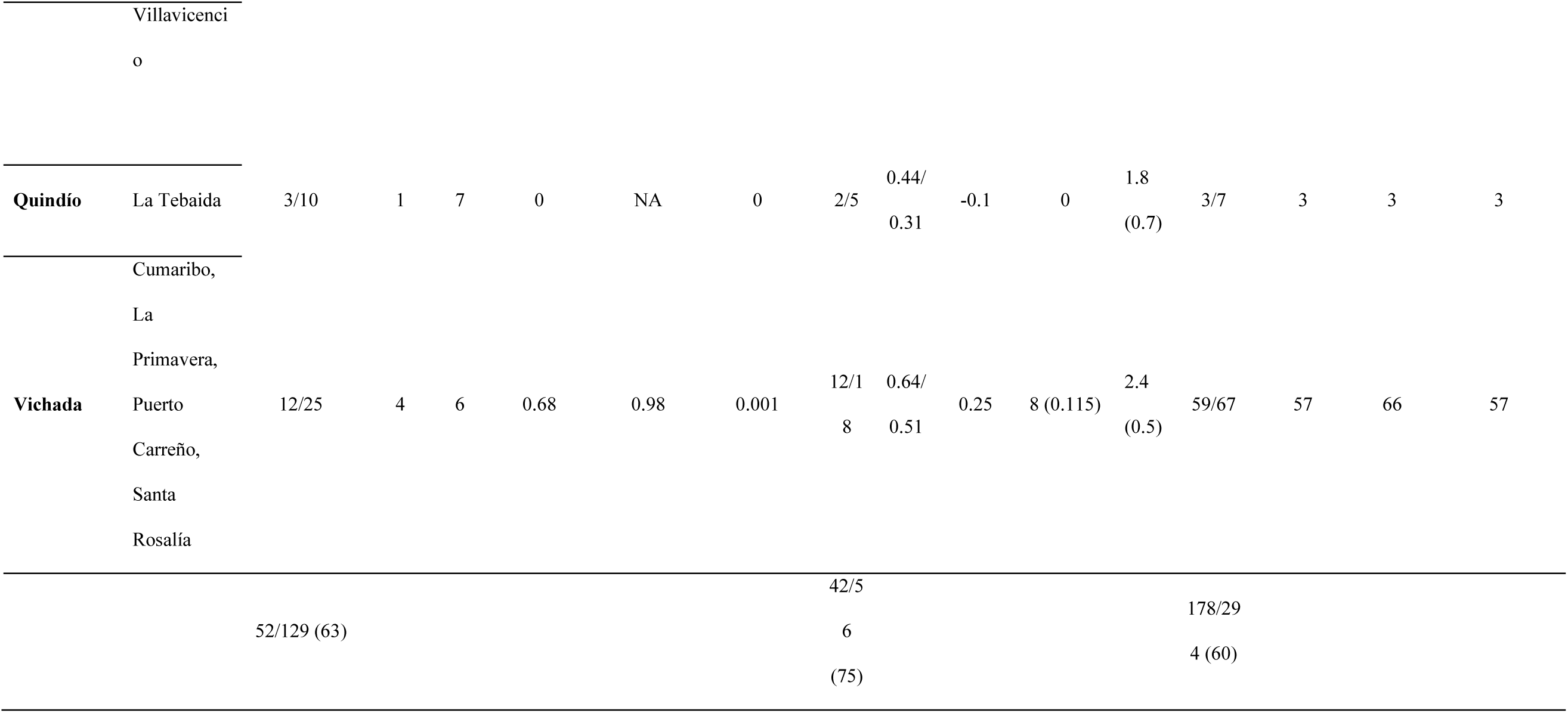
Summary of arbovirus natural infection, genetic characterization, and Wolbachia screening for *Aedes albopictus* populations from Colombia.

Larvae were collected from a variety of artificial and natural breeding habitats as described by [5], preserved in absolute ethanol or RNAlater, and taxonomically identified in each department using standard morphological criteria as described previously [22,23]. All specimens were subsequently transferred to Bogotá for confirmation and staging under a Zeiss Stemi DV4 stereomicroscope (Zeiss, Jena, Germany). Only fourth instar (L4) larvae were retained for downstream molecular analyses and processed individually to ensure optimal DNA yield and integrity.

### DNA and RNA extraction

To minimize pseudo replication, only one larva per breeding site was analyzed, and sites separated by ≥ 500 m were included, following best practices recommendations for *Aedes albopictus* population-genetic studies [24]. Each larva preserved in RNAlater was transferred to 200 µL of diethyl-pyrocarbonate-treated water (Sigma-Aldrich D5758) and manually homogenized using sterile disposable pestles (USA Scientific 1415-5390). Simultaneous extraction of DNA and RNA was performed using the Viral Nucleic Acid Extraction Kit II (IBI Scientific, Dubuque, USA) according to the manufacturer’s protocol, with the lysis-buffer incubation extended to 1 h to maximize nucleic acid recovery. Extracted nucleic acids were quantified using a NanoPhotometer NP80 (Implen, California, USA).

### Mitochondrial COI haplotype analyses

Two overlapping fragments of the COI gene were amplified with primers pairs 1454F/2160R and 2027F/2886R following published PCR protocol by [10]. Amplicons were verified on 1% agarose gels stained with ethidium bromide and subsequently sequenced bidirectionally via capillary electrophoresis on and ABI 3730xl platform. Forward and reverse chromatograms were assembled into consensus sequences (∼1350 bp) using DNASTAR Lasergene, verified by BLAST, searches and aligned against reference sequences from GenBank database using MEGA and AliView. Haplotype inference, and networks were reconstructed in PopART using the median-joining algorithm, to facilitate visualization we grouped Colombian samples as departments and international references sequences as Countries. Mitochondrial diversity indices including the number of segregating sites (S), haplotypes (h), haplotype diversity (Hd), nucleotide diversity (π), mean number of pairwise differences (k), and Tajima’s D values were estimated using the *ape* and pegas packages in R. Population differentiation was assessed using ΦST values under the K80 substitution model with 1,000 permutations. Bayesian phylogenetic inference was performed in MrBayes, and the resulting trees and haplotype networks were visualized and annotated for interpretation. All COI sequences generated in this study were deposited in GenBank under accession numbers P Z137697 –PZ137748 (Data File S1).

### Microsatellite genotyping and analyses

Individuals were genotyped using 11 microsatellite loci previously validated for *Ae. albopictus* populations analyses [7,12]. DNA samples were diluted to 7 ng/µL in nuclease free water prior to PCR amplification, Microsatellite loci (Aealbmic1, Aealbmic2, Aealbmic3, Aealbmic5, Aealbmic6, Aealbmic9, Aealbmic11, Aealbmic14, Aealbmic15, Aealbmic16, and Aealbmic17) were amplified using primer sequence and protocols described by (Manni et al., 2015). Amplicons were checked by 2 % agarose electrophoresis and genotyped on an ABI3730xl DNA Analyzer (Applied Biosystems). Allele binning was automated with TANDEM v1.09 and manually curated to resolve ambiguous calls. Failed or ambiguous amplification were re-amplified and genotyped for the affected locus.

The final microsatellite dataset included Colombian populations together with reference multilocus genotypes from additional geographic regions obtained from[7]. Locus quality control included inspection of allele-frequency distributions, missing data patterns, Hardy–Weinberg equilibrium deviations, and locus-specific differentiation metrics. Three loci (Aealbmic15, Aealbmic6, and Aealbmic3) were excluded following quality-control analyses because they showed inconsistent amplification performance, atypical allele-frequency distributions, elevated missing-data rates, and disproportionate effects on estimates of global population structure. Subsequent analyses were therefore conducted using the remaining eight loci that exhibited more stable genotyping performance across populations, resulting in a final dataset of eight polymorphic microsatellite loci (Aealbmic1, Aealbmic2, Aealbmic5, Aealbmic9, Aealbmic11, Aealbmic14, Aealbmic16, and Aealbmic17).

Genetic diversity indices, including observed heterozygosity (HO), expected heterozygosity (HE), and inbreeding coefficients (FIS), were estimated using hierfstat [25]. Deviations from Hardy–Weinberg equilibrium were evaluated for each locus using pegas. Global and pairwise population differentiation were estimated using Weir and Cockerham’s FST, and visualized using heatmaps generated with pheatmap [26].

Multivariate genetic structure was explored using Principal Coordinates Analysis (PCoA) based on pairwise genetic distances and Discriminant Analysis of Principal Components (DAPC) implemented in adegenet [27]. The number of retained principal components was optimized by cross-validation, and alternative clustering solutions were evaluated across multiple K values using Bayesian Information Criterion (BIC) profiles. Because the K = 8 configuration captured the major components of global admixture and improved geographic interpretability, this configuration was retained for downstream visualization and comparative analyses. DAPC membership probabilities were additionally projected onto global geographic maps to visualize spatial patterns of ancestry composition and potential connectivity among populations.

Hierarchical genetic partitioning was evaluated through Analysis of Molecular Variance (AMOVA) implemented in poppr using a region/population hierarchical design and 999 permutations. Potential demographic instability was evaluated using the Garza–Williamson M-ratio [28]. All analyses were conducted in R v4.3.3, with data handling and visualization performed using dplyr, ggplot2, adegenet, poppr, ape, scatterpie, and related packages.

National connectivity analysis:

National-scale transport connectivity among Colombian departments was evaluated using terrestrial cargo mobility data obtained from the web page of the Colombian Ministry of Transport (https://plc.mintransporte.gov.co/Estad%C3%ADsticas/Carga-Modo-Terrestre/Carga-mensual-RNDC). Transport variables included the cumulative number of cargo trips and transported tonnage between sampled departments during the study period. Pairwise road distance and travel time estimates between departments were additionally incorporated as proxies of geographic connectivity. Transport metrics were log10-transformed prior to analysis to improve comparability among routes. Pairwise genetic differentiation matrices derived from microsatellite (linearized FST) and mitochondrial COI (ΦST–K80) datasets were independently evaluated against geographic and transport connectivity matrices using Mantel and partial Mantel tests implemented in the vegan package in Rv4.3.3. Partial Mantel tests were used to assess associations between mitochondrial and microsatellite differentiation and transport connectivity while controlling for geographic distance.

### Statistical analyses linking genetic differentiation and trade flows

To evaluate whether international trade influenced genetic differentiation among *Ae. albopictus* populations, pairwise microsatellite differentiation (FST) and mitochondrial differentiation (ΦST–K80 from COI sequences) were analyzed separately using regression and correlation-based approaches. Colombian samples were grouped by department, whereas international reference populations were defined according to the most specific locality available for each marker dataset. Pairwise genetic matrices were integrated with geographic distance and trade-associated predictors derived from Colombian customs records (DIAN), available at https://www.dian.gov.co/dian/cifras/Paginas/Bases-Estadisticas-de-Comercio-Exterior-Importaciones-y-Exportaciones.aspx, including import tonnage and shipment frequency for high-risk commodities (used tires, new tires, and live plants), as well as combined “total risk” metrics integrating these categories for country of origin and reported destination Colombian department. Trade variables were log10-transformed prior to analysis.

For microsatellite analyses, pairwise FST values between Colombian departments and international populations were modeled as a function of geographic distance and trade intensity using linear models implemented in base R. Competing models were compared using Akaike Information Criterion (AIC), adjusted R², and overall model significance. Correlation analyses were additionally performed using Spearman’s rank tests. To evaluate potential spatial heterogeneity among Colombian regions, department-specific associations between geographic distance, trade variables, and genetic differentiation were explored independently. A similar analytical workflow was applied to mitochondrial COI differentiation (ΦST–K80). Variance inflation factors (VIFs) were evaluated to assess multicollinearity among predictors. Residual diagnostics included visual inspection of fitted-versus-residual plots, Cook’s distance, leverage statistics, and Shapiro–Wilk tests of residual normality. Regression analyses, model comparisons, and correlation tests were conducted in R v4.3.3.

Sensitivity analyses were conducted to evaluate model robustness. A leave-one-department-out approach was applied by iteratively excluding each Colombian department and refitting linear models relating microsatellite differentiation (FST) to geographic distance and total-risk shipment frequency using functions from the stats, dplyr, purrr, and broom packages in R. Model coefficients, adjusted R², and AIC values were compared across iterations. Permutation analyses were additionally performed by randomly permuting trade predictors and comparing observed coefficients and R² values against null distributions generated from 9,999 permutations using stats and ggplot2. Multicollinearity was evaluated using VIF/GVIF metrics implemented in the car package, and residual diagnostics were assessed through inspection of residual distributions, leverage, and Cook’s distance values using base R functions.

To visualize potential introduction pathways, pairwise microsatellite similarity (1 − FST) was integrated with international trade connectivity. Spatial networks and regression figures were generated using ggplot2, ggpubr, patchwork, sf, and maps-based visualization workflows.

### Wolbachia screening and prevalence analyses

Detection of *Wolbachia* infection was performed by PCR amplification of the 16S rRNA and wsp genes. Samples yielding both amplicons were considered positive. Positive samples were further screened using lineage-specific primers for wAlbA and wAlbB to determine infection types (A, B, or A+B). Prevalence was calculated at the department level with 95% binomial confidence intervals. Spatial patterns of infection were visualized using the sf package, rnaturalearth, and scatterpie. To explore potential associations between *Wolbachia* infection dynamics and mito-nuclear population structure, *Wolbachia* prevalence (total infection, wAlbA, wAlbB, and A+B coinfection frequencies) was compared with mitochondrial (haplotype diversity (Hd), nucleotide diversity (π), and pairwise ΦST estimates derived from COI sequences) and nuclear genetic (microsatellite-based FST estimates) diversity and differentiation indices across Colombian populations of *Ae. albopictus*. Mito-nuclear discordance was evaluated as the difference between pairwise mitochondrial and nuclear differentiation values, Φ_*ST*_ − *F*_*ST*_, , where positive values indicate proportionally greater mitochondrial differentiation relative to nuclear structure. Pairwise differences in Wolbachia prevalence and coinfection frequencies between populations were subsequently compared against mitochondrial differentiation, nuclear differentiation, and mito-nuclear discordance metrics using Spearman’s rank correlations and exploratory linear regression models implemented in R v4.3.3.

### Arbovirus detection

Viral RNA was reverse-transcribed and amplified using the Luna Universal One-Step RT-PCR kit (Thermo Fisher Scientific, USA) and multiplex nested RT-PCR system targeting DENV, CHIKV, and ZIKV as previously described [29]. RNA extracted from culture-harvested viral stocks was used as positive controls, whereas nuclease-free water served as the negative control. PCR amplicons were separated on 2 % agarose gels stained with ethidium bromide and visualized under ultraviolet illumination to confirm virus-specific amplification profiles.

## RESULTS

A total of 657 *Ae. albopictus* larvae were collected from five Colombian departments: Antioquia (8.0%), Cauca (43.7%), Meta (23.9%), Quindío (1.5%), and Vichada (22.9%) (Table 1). Sampling encompassed 18 municipalities, including one in Antioquia, eight in Cauca, six in Meta, one in Quindío, and four in Vichada, representing the Andean, Pacific, and Orinoquia regions of Colombia.

### Mitochondrial COI haplotype analyses

A total of 129 *Ae. albopictus* larvae from different localities were processed for COI amplification. Dual fragment amplicons were successfully obtained from 52 Colombian specimens, with sequences distributed as follows: Cauca (48%), Vichada (23%), Meta (15%), Antioquia (8%), and Quindío (6%) (Table 1). To provide a broader geographic context, 88 additional COI sequences were downloaded from GenBank data base, resulting in a final dataset of 140 sequences representing 82 haplotypes.

Overall, the dataset exhibited low nucleotide diversity (π = 0.002) and very high haplotype diversity (Hd = 0.95). We identified 42 segregating sites, of which 30 were parsimony-informative, indicating sufficient phylogenetic signal for the construction of a stable haplotype network. Tajima’s D value for the global dataset was negative (D = –1.45, p = 0.93), although not statistically significant, suggesting no strong departure from neutrality. Across Colombian populations, COI sequences also showed consistently low nucleotide diversity (π ≈ 0.0004–0.0025) alongside moderate-to-high haplotype diversity (Hd ≈ 0.48–0.70). Tajima’s D values were negative in Antioquia, Cauca and Meta, indicating an excess of low frequency variants consistent with recent expansion, although the small sample sizes limit interpretive power. In contrast, Vichada displayed a positive Tajima’s D value, which may reflect potential population structure or admixture (Table 1).

Pairwise ΦST values revealed low but heterogeneous mitochondrial differentiation both within Colombia and relative to international populations, indicating contrasting patterns of mitochondrial connectivity among departments. Antioquia and Meta showed comparatively lower differentiation from USA populations (ΦST = 0.22 and 0.08, respectively), particularly with Los Angeles populations, while Meta and Vichada also exhibited moderate differentiation from East and Southeast Asian regions, including Singapore (ΦST = 0.08–0.22) and China/Taiwan/Japan (ΦST = 0.21–0.27), with zero differentiation detected between Meta and populations from Los Angeles (USA) and Rio de Janeiro (Brazil) (Figure S1; Table S1). In contrast, Cauca and Quindío displayed markedly higher divergence from Chinese populations (e.g., Cauca–China ΦST = 0.59**, Quindío–China ΦST = 0.64**), despite showing affinity with Singapore populations and, in the case of Quindío, with Meta.

Consistent with these patterns, the median-joining haplotype network recovered 82 haplotypes, of which 67 (81.7%) were country-specific and only 15 were shared among countries (Figure 2A). Three main mitochondrial clusters were identified, with the largest centered on a widespread haplotype (Hap_18) shared by Antioquia, Meta, Quindío, Cauca, and Singapore (Figure 2B). Several Cauca-specific haplotypes radiated from this lineage, consistent with local diversification around a shared ancestral haplotype. In contrast, Vichada haplotypes were mostly short terminal branches associated with internationally distributed lineages. For example, Hap_16 was separated by a single mutational step from a Malaysian-associated haplotype (Hap_1) also connected to Brazil, Cauca, and USA sequences, while Hap_14 was shared with Meta, Antioquia, Italy, Mexico, and Japan and showed no associated local derivatives (Figure 2C). Together, these mitochondrial patterns are consistent with multiple lineage affinities across Colombian departments, with some populations showing evidence of local haplotype diversification, whereas others remained closely associated with widely distributed international lineages.

**Figure 2.**
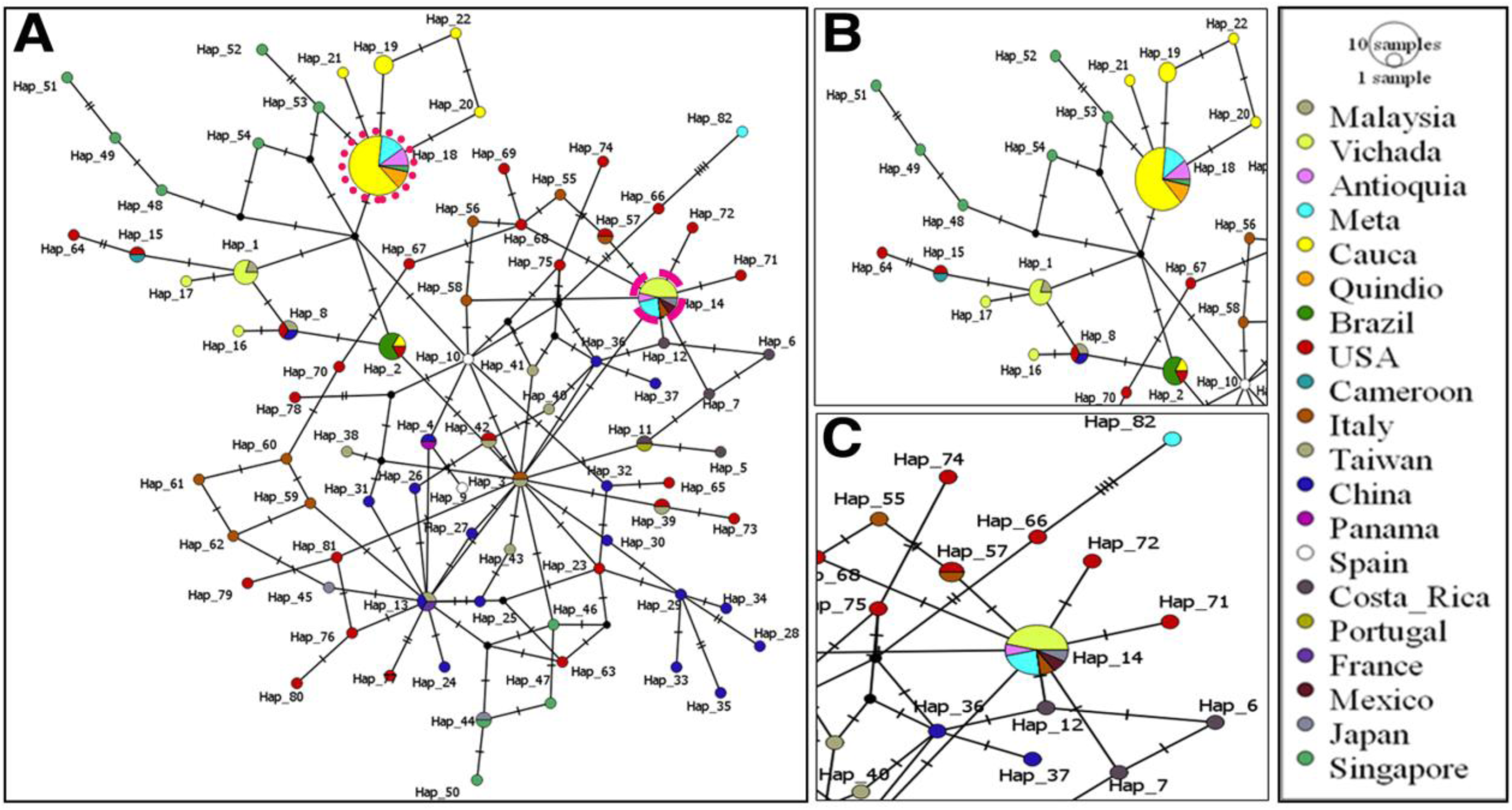
Mitochondrial haplotype structure of *Aedes albopictus* inferred from COI sequences. Median-joining haplotype networks were reconstructed from a 1,350-bp mitochondrial COI alignment including Colombian and international reference sequences (n = 140). Each node represents a unique haplotype (Hap_x), with node size proportional to haplotype frequency. Pie charts indicate the geographic origin of sequences composing each haplotype according to the legend. Lines between nodes represent mutational steps, with hatch marks indicating nucleotide substitutions; small black nodes correspond to inferred median vectors (unsampled or intermediate haplotypes). (A) Global haplotype network showing overall mitochondrial relationships among Colombian and international populations. Three major mitochondrial clusters were identified, including a widespread central lineage shared across multiple geographic regions. (B) Magnified view of the cluster centered on Hap_18, a common haplotype shared among several Colombian departments and Singapore, from which multiple Cauca-associated haplotypes radiate, suggesting local diversification following introduction. (C) Magnified view of the Hap_14 cluster, showing haplotypes shared between Vichada, Meta, Antioquia, and international populations from Italy, Mexico, and Japan, consistent with recent and geographically widespread introduction lineages.

The haplotype network, dominated by closely connected haplotypes with few mutational steps separating them, suggested very shallow evolutionary divergence. To formally assess whether this pattern was also reflected at deeper branching levels, we inferred Bayesian phylogeny (Figure S 2). Only a few shallow clades achieved moderate support, while the backbone remained largely unresolved. This unresolved topology is fully consistent with the demographic signals from the diversity metrics and the network analysis, all pointing to a recent, demographic expansion with limited phylogenetic structure despite high haplotype richness.

### Microsatellite analysis

Microsatellite loci successfully amplified in 75% of processed individuals. Preliminary quality-control analyses identified three loci (Albmic15, Albmic6, and Albmic3) showing inconsistent amplification performance, atypical allele-frequency distributions, elevated missing-data rates, and disproportionate effects on estimates of global genetic differentiation. The mean FST across all loci was 0.156; exclusion of these loci reduced the estimate to 0.058. Consequently, subsequent analyses were conducted using the remaining eight loci. After filtering, microsatellite data revealed moderate but significant population structure. AMOVA results indicated that 7.03% of variance was explained by differences between Colombian and external populations, 16.49% by differences among populations within groups, and 76.48% by variation among individuals (all p < 0.005).

Pairwise microsatellite FST values revealed distinct nuclear structuring among Colombian departments (Figure S3; Table S2). Quindío exhibited the highest divergence and emerged as the most genetically distinctive profile. Cauca displayed intermediate differentiation, reflecting partial isolation. In contrast, Meta and Vichada exhibited comparatively low differentiation from one another, consistent with either recent connectivity or shared ancestry. Comparing Colombian populations with other countries, Colombian were genetically closer to USA and several East–Southeast Asian references, and more differentiated from many Latin American populations (e.g., Panama, Argentina, Brazil), but strong influence of France, La Reunion, Hawaii, And Japan. This pattern suggests that Colombian populations share genetic affinities with multiple international regions rather than exclusively with neighboring Latin American populations, consistent with a complex invasion history involving geographically diverse lineages (Figure S3).

DAPC analyses were interpreted using the K = 8 clustering solution, which corresponded to the region where BIC improvement began to plateau (Figure S4A) and provided biologically interpretable patterns of geographic structure, this solution was also broadly consistent with previous analyses based on the same microsatellite panel [6]. Under this clustering solution, Colombian populations formed a partially differentiated assemblage while exhibiting varying levels of membership in clusters shared with Asian, European, and Central American populations (Figure 3A, Figure 4BS). The analysis reveals a clear genetic structure, with the Colombian populations (Meta, Antioquia, Cauca, Quindío, Vichada) showing partial differentiation relative to international reference samples by PCoA analysis (Figure S5). This suggests both regional divergence and shared ancestry across continents. Despite this mixed ancestry, a consistent trend emerged: cluster 6 was dominant across all Colombian populations (figure S4B). This cluster includes populations from Mexico, Greece, and the USA. Patterns of secondary membership varied among departments: Cauca showed contributions from clusters 1 (31%) and 4 (21.3%). Meta and Vichada displayed minor membership in cluster 1 (17% and 25% respectively), and Quindío showed 26% assignment to cluster 2 (Figure S4B). Overall, these results indicate that Colombian *Ae. albopictu*s populations share a common dominant genetic component while also exhibiting heterogeneous admixture patterns associated with multiple international genetic affinities.

**Figure 3.**
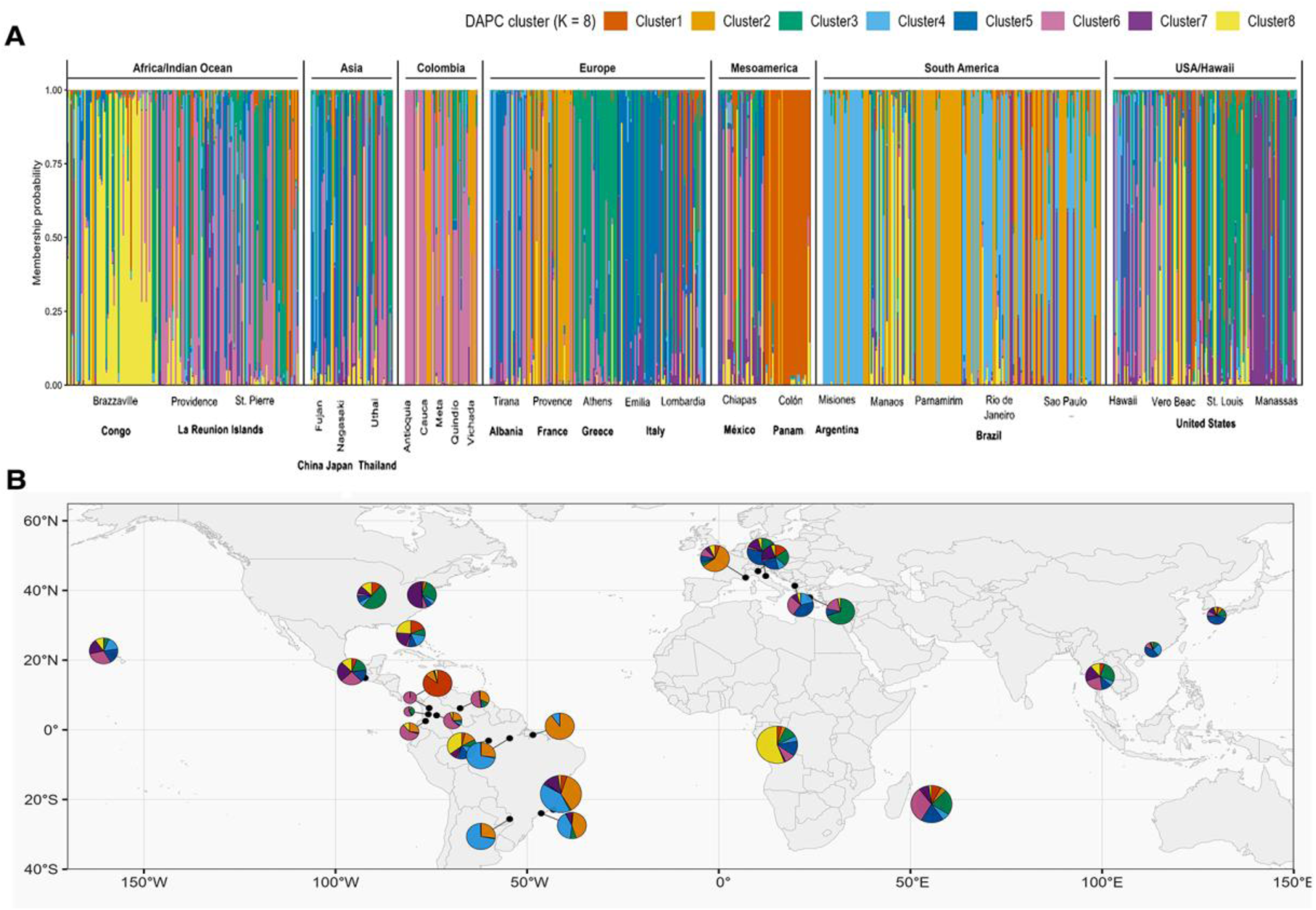
Global nuclear genetic structure of *Aedes albopictus* inferred from microsatellite data. (A) Discriminant Analysis of Principal Components (DAPC) membership probabilities (K = 8) for individual mosquitoes grouped by geographic region and sampling locality. Each vertical bar represents one individual, and colors indicate proportional assignment to inferred nuclear genetic clusters. Colombian populations exhibit mixed ancestry profiles shared with populations from Asia, Europe, the Americas, and Indian Ocean regions, consistent with multiple introduction sources and extensive admixture. (B) Geographic distribution of DAPC cluster composition across sampled localities worldwide. Pie charts represent the proportional frequency of inferred nuclear clusters at each sampling site. Colombian populations display heterogeneous cluster composition, with some departments sharing ancestry components with Asian, European, and American populations, supporting complex invasion pathways and repeated introductions.

**Figure 4.**
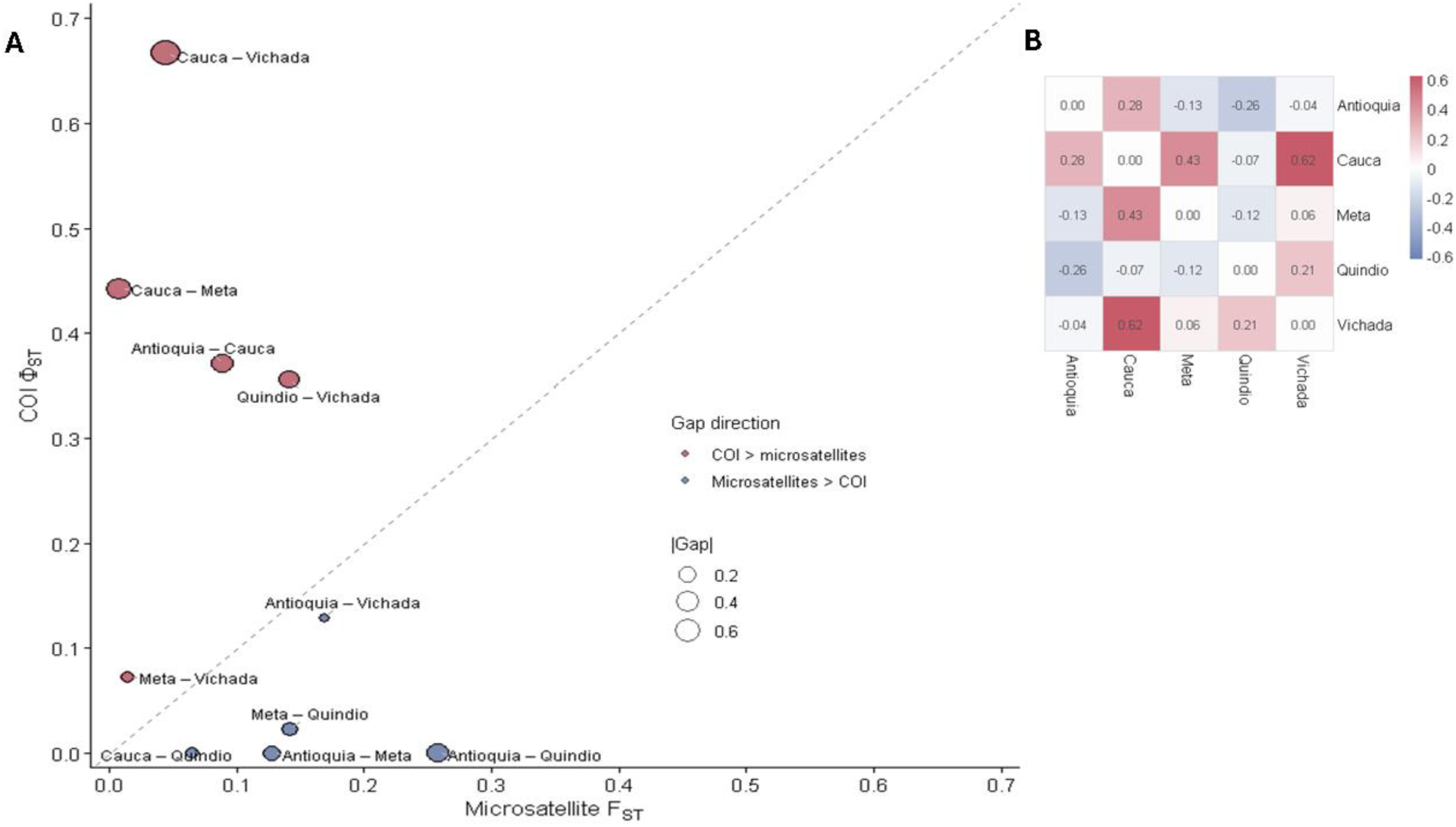
Mito–nuclear differentiation gap among Colombian *Aedes albopictus* populations. (A) Pairwise comparison between mitochondrial differentiation (COI ΦST–K80) and nuclear differentiation (microsatellite FST) among Colombian departments. Each point represents a pairwise population comparison, with point size proportional to the absolute magnitude of the mito–nuclear differentiation gap. The dashed diagonal indicates equality between mitochondrial and nuclear differentiation. Red points indicate comparisons in which mitochondrial differentiation exceeded nuclear differentiation, whereas blue points indicate the opposite pattern. Several population pairs, particularly involving Cauca and Vichada, showed substantially higher mitochondrial than nuclear differentiation, consistent with mito–nuclear spatial decoupling. (B) Heatmap showing the signed pairwise differentiation gap calculated as COI ΦST minus microsatellite FST. Positive values (red) indicate greater mitochondrial differentiation relative to nuclear differentiation, whereas negative values (blue) indicate relatively stronger nuclear differentiation. Overall, the observed asymmetry supports contrasting spatial dynamics between maternally inherited mitochondrial lineages and recombining nuclear markers across Colombian populations.

The Garza–Williamson M-ratio was consistently low across populations (0.33–0.65), remaining below the commonly used threshold of 0.68 and suggesting recent demographic instability compatible with bottlenecks, founder effects, or other processes associated with invasion dynamics. Taken together, these analyses reveal heterogeneous nuclear genetic structure among Colombian *Ae. albopictus* populations, characterized by admixture, partial differentiation, and signatures consistent with recent demographic instability.

### Mito-nuclear genetic decoupling

Mito-nuclear comparisons revealed heterogeneous patterns of discordance among Colombian *Ae. albopictus* populations (Fig 4). Pairwise comparisons between nuclear differentiation (FST) and mitochondrial differentiation (ΦST) showed that several population pairs deviated markedly from the expected 1:1 concordance pattern, indicating that nuclear and mitochondrial markers captured partially distinct patterns of population differentiation (Fig 4A). The strongest mito-nuclear gaps were observed for comparisons involving Cauca, particularly Cauca–Vichada and Cauca–Meta, where mitochondrial differentiation remained high despite low nuclear differentiation. In contrast, other comparisons, such as Antioquia–Quindío, showed relatively higher nuclear differentiation but limited mitochondrial divergence. Heatmap visualization of the mito-nuclear gap (ΦST − FST) further demonstrated that discordance was not uniformly distributed across populations but instead showed directional and population-specific patterns (Fig 4B). Overall, these results suggest that nuclear and mitochondrial markers captured partially distinct spatial patterns among Colombian populations. Whereas nuclear differentiation was generally lower among several population pairs, mitochondrial differentiation remained comparatively elevated in specific comparisons, consistent with heterogeneous mito-nuclear structure across the Colombian invasion landscape.

### National Scale analysis of geographic distance and commercial connectivity

National-scale analyses revealed contrasting spatial patterns between mitochondrial and nuclear markers (Figure 5; Table S3). Mitochondrial differentiation based on COI sequences showed a strong and significant positive association with geographic distance (Mantel r = 0.79, p = 0.033), indicating partial isolation by distance among Colombian populations of *Ae. albopictus.* In contrast, microsatellite differentiation showed no significant association with geographic distance (Mantel r = 0.15, p = 0.350), suggesting limited nuclear spatial structuring at the national scale.

**Figure 5.**
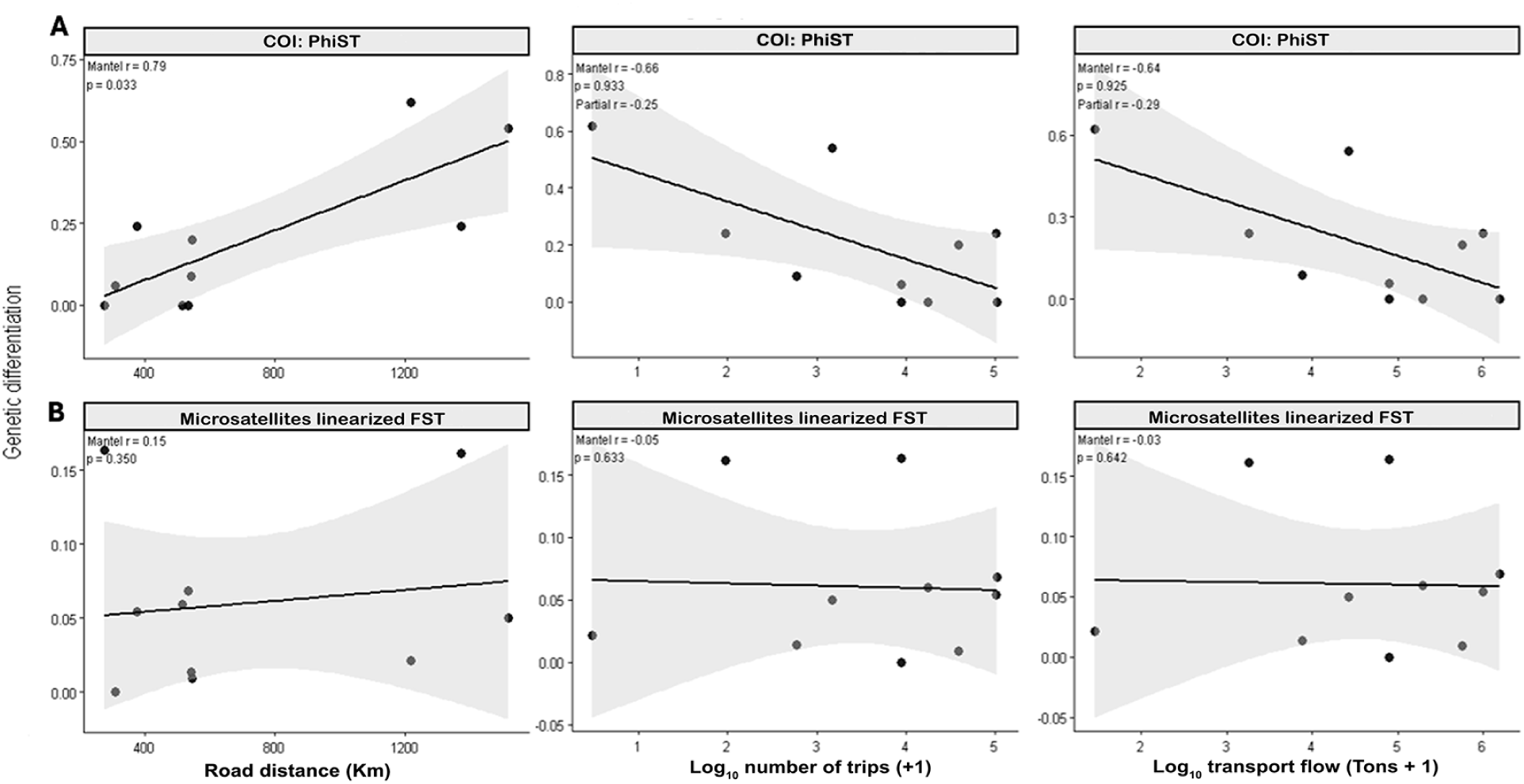
National-scale geographic and transport predictors of genetic differentiation in Colombian *Aedes albopictus* populations. Relationships between pairwise genetic differentiation and geographic or transport connectivity metrics among Colombian departments. Upper panels (A) show mitochondrial differentiation based on COI ΦST–K80 values, whereas lower panels (B) show nuclear differentiation based on linearized microsatellite FST values. Genetic differentiation was evaluated against pairwise road distance, log10-transformed number of cargo trips, and log10-transformed transported tonnage derived from the Colombian National Freight Dispatch Registry (RNDC). Solid lines represent linear regression fits with 95% confidence intervals. Mantel correlation coefficients and associated permutation-based p-values are shown within each panel; partial Mantel statistics controlling for geographic distance are additionally indicated for transport connectivity analyses involving mitochondrial differentiation. Mitochondrial differentiation showed a significant positive association with geographic distance, whereas microsatellite differentiation exhibited weak and non-significant spatial structure.

Mitochondrial differentiation additionally showed moderate negative associations with transport connectivity proxies, including cargo flow tonnage (Mantel r = –0.64, p = 0.925) and number of trips (Mantel r = –0.66, p = 0.933), although neither relationship was statistically significant.

The contrasting Mantel results reinforce the mito-nuclear discordance identified in previous analyses, with mitochondrial differentiation retaining a detectable geographic signal, whereas contemporary nuclear structure appeared largely decoupled from geographic distance. Together, these patterns suggest that mitochondrial and nuclear markers capture different spatial dimensions of population structure within Colombia.

Eco-logistic profiling of sampled departments further revealed substantial environmental and anthropogenic heterogeneity across the Colombian invasion landscape (Figure S6). The sampled populations occupied environmentally distinct settings that differed in climatic conditions, urbanization intensity, and human mobility characteristics, providing ecological context for the observed patterns of genetic differentiation.

### Association between international trade and genetic differentiation

Association between international trade and genetic differentiation analyses based on nuclear microsatellite data revealed consistent relationships between trade connectivity, commodity type, and population differentiation among Colombian *Ae. albopictus* populations. Geographic distance alone was negatively associated with pairwise microsatellite FST (R² = 0.15, adjusted R² = 0.14, p < 0.001; Figure 6A, Table 2). However, models incorporating trade-associated predictors substantially improved explanatory power. The strongest model included total-risk shipment frequency, integrating used tires, new tires, and live plants, explaining 34.1% of the variance in microsatellite differentiation (adjusted R² = 0.33; AIC = −366.36; p < 0.001). Models based on used tire shipment frequency also showed strong support (adjusted R² = 0.28; AIC = −358.64; p < 0.001). Interaction models incorporating department-specific geographic effects further improved model fit (adjusted R² = 0.54; AIC = −404.08; p < 0.001), indicating heterogeneous geographic–genetic relationships among Colombian departments. Similar trends were observed when analyses were performed using import tonnage rather than shipment frequency (Figure S7). Across analyses, higher trade connectivity was consistently associated with higher microsatellite differentiation values (Figure 6B–C; Figure S7; Table 2).

**Figure 6.**
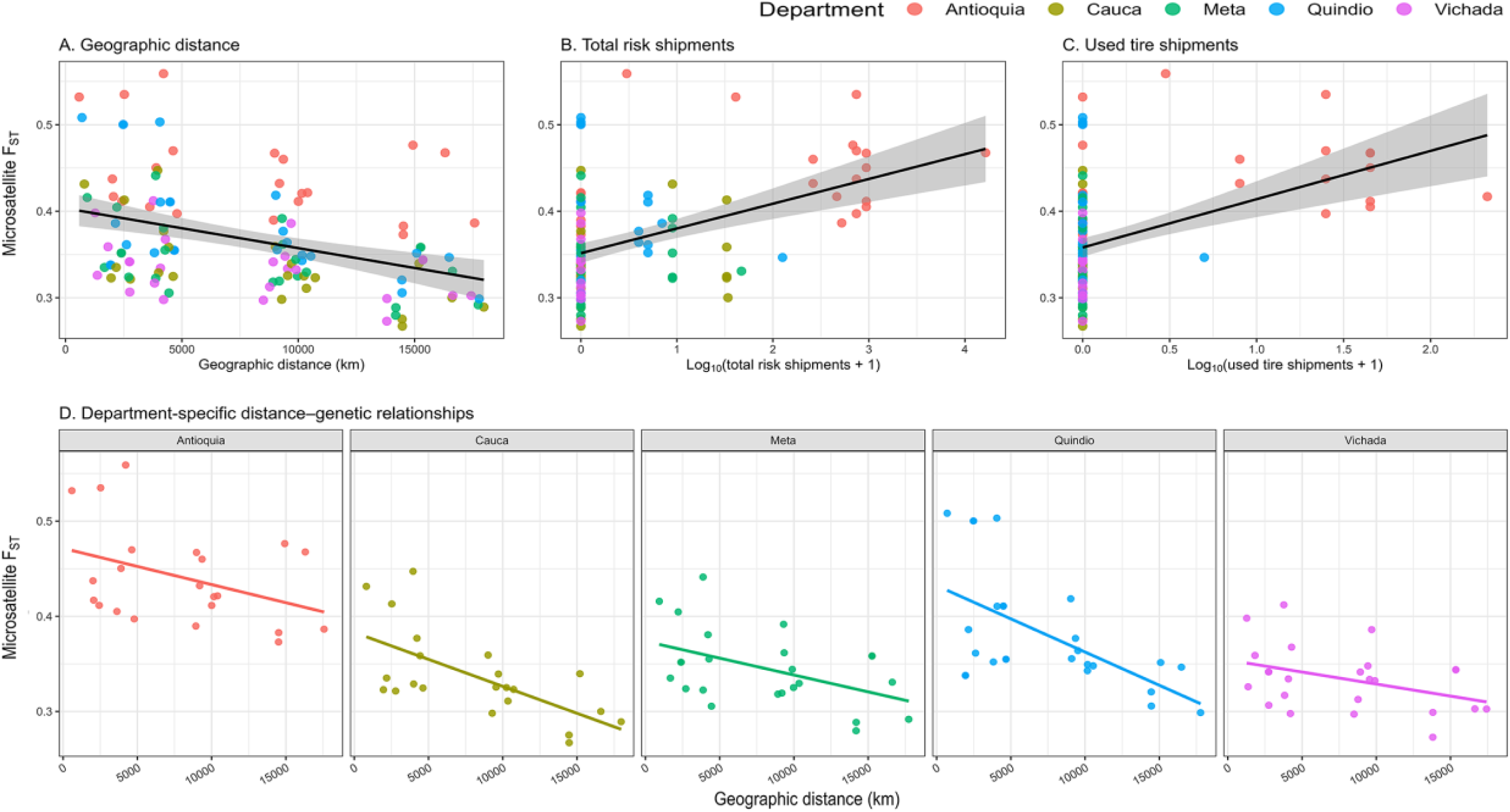
International trade and geographic distance associated with microsatellite differentiation in Colombian *Aedes albopictus* populations. (A) Relationship between pairwise microsatellite differentiation (FST) and geographic distance between Colombian departments and international *Aedes albopictus* populations. (B) Association between microsatellite differentiation and total-risk shipment frequency, integrating used tires, new tires, and live plants. (C) Association between microsatellite differentiation and shipment frequency of used tires. Solid lines represent linear regression fits with 95% confidence intervals shaded in gray. Points are colored according to Colombian department. (D) Department-specific relationships between geographic distance and microsatellite differentiation for Antioquia, Cauca, Meta, Quindío, and Vichada, illustrating heterogeneous geographic–genetic patterns among departments. Trade-associated predictors and geographic distance were log10-transformed prior to analysis.

**Table 2.**
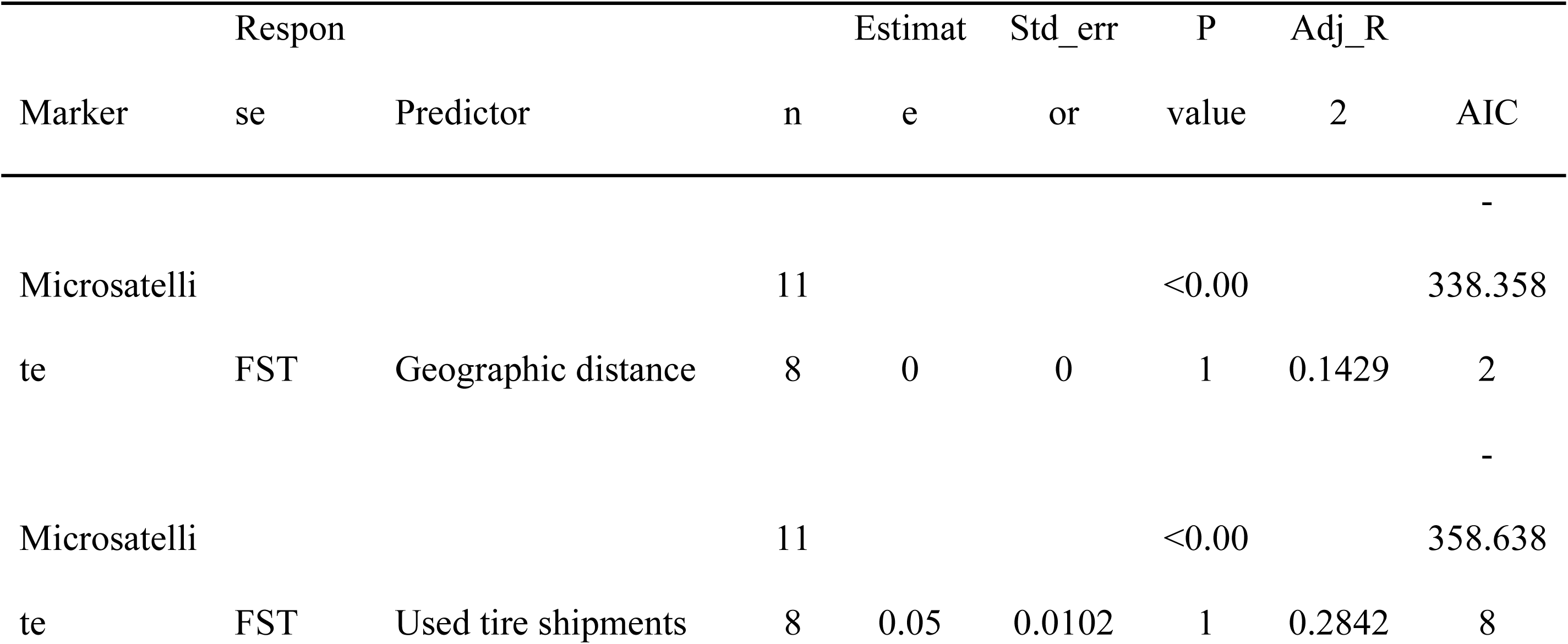
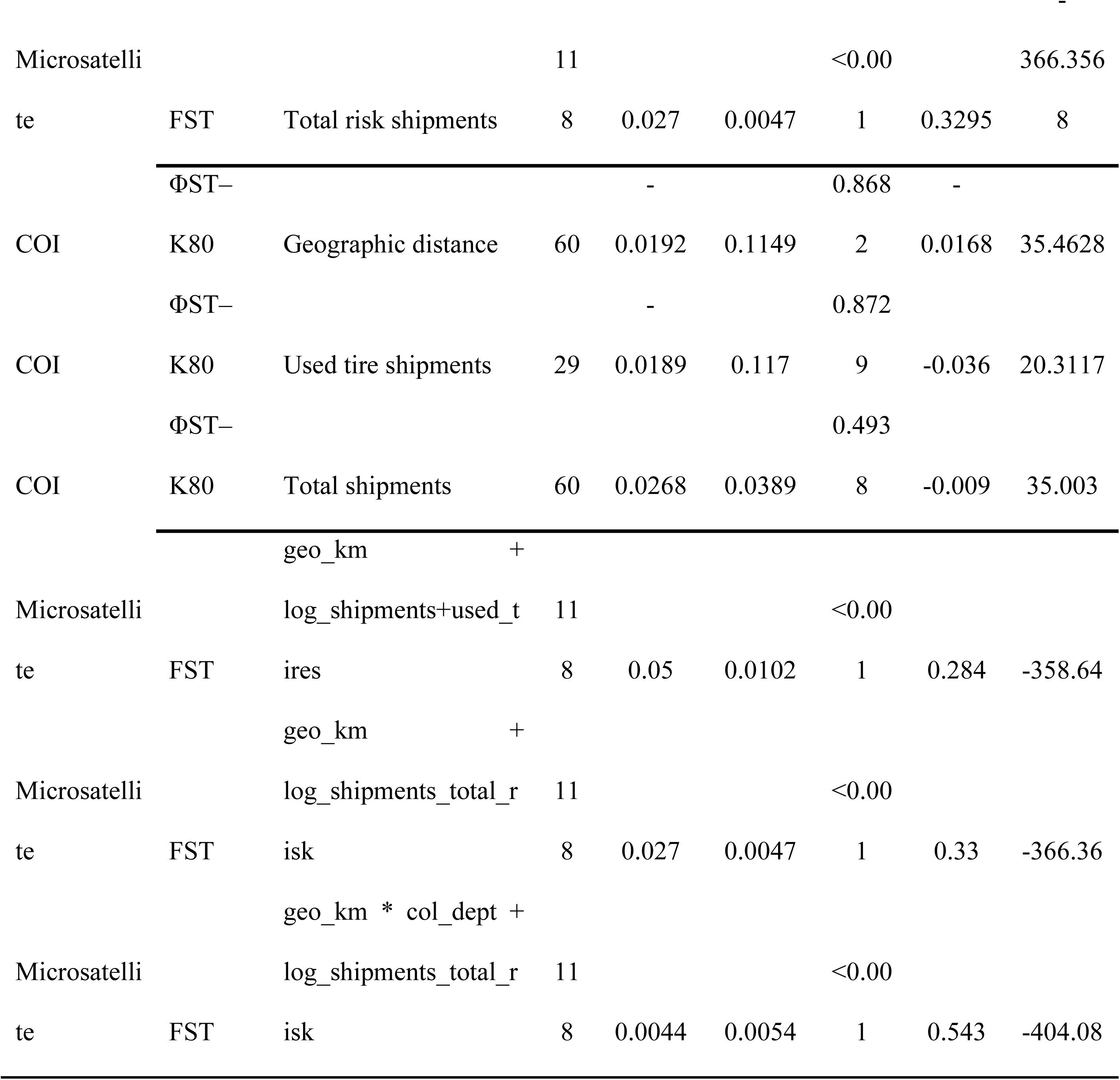
Linear regression models evaluating associations between geographic distance, international trade connectivity, and genetic differentiation in Colombian *Aedes albopictus* populations using microsatellite and mitochondrial COI markers.

Correlation analyses supported these patterns. Total-risk import tonnage (Spearman’s ρ = 0.438, p < 0.001), used tire tonnage (ρ = 0.437, p < 0.001), total-risk shipment frequency (ρ = 0.434, p < 0.001), and used tire shipment frequency (ρ = 0.434, p < 0.001) showed the strongest associations with microsatellite FST. Comparable but generally weaker relationships were detected for Jost’s D and GST′ (Table S4). Live plant-associated variables exhibited lower effect sizes but remained significant, whereas new tire-associated variables also showed positive associations with differentiation, particularly for shipment frequency metrics (Figure S7).

Department-specific analyses revealed heterogeneous trade-associated differentiation patterns among Colombian regions (Figure 6D). Antioquia consistently exhibited comparatively higher microsatellite FST values across international comparisons and stronger relative associations with used tire-related trade variables, whereas Cauca, Meta, and Quindío showed comparatively stronger associations with total-risk and new tire shipment metrics. Live plant-associated relationships were generally weaker and inconsistent among departments. Sensitivity analyses demonstrated that the observed trade-associated signal was robust to permutation-based null expectations (empirical p < 0.0001). Leave-one-department-out analyses further showed that positive trade effects remained generally consistent across departments, although exclusion of Antioquia substantially reduced model fit and effect size, indicating that this department accounted for a substantial portion of the overall trade-associated differentiation signal.

In contrast, mitochondrial COI differentiation exhibited weaker and non-significant associations with trade variables (Figure S8; Table 2). Geographic distance showed minimal association with ΦST–K80, and shipment-related predictors explained substantially less variance than observed for microsatellite markers. Department-level mitochondrial patterns were also heterogeneous, with Cauca and Quindío displaying comparatively elevated ΦST values across international comparisons, whereas Antioquia, Meta, and Vichada showed weaker or more variable relationships. Together, these results indicate that trade-related connectivity patterns are more strongly reflected in contemporary nuclear microsatellite structure than in mitochondrial variation. This contrast reinforces the mito-nuclear discordance identified in previous analyses and suggests that contemporary trade-associated connectivity is more closely associated with nuclear genetic differentiation than with mitochondrial lineage structure.

### Trade-genetic connectivity patterns among Colombian and international populations

To visualize the overlap between international trade connectivity and nuclear genetic similarity, we integrated microsatellite differentiation metrics with trade-flow information among Colombian departments and international populations. The combined analysis of genetic similarity and international trade connectivity identified multiple geographically plausible trade-genetic connectivity patterns linking Colombian departments with international *Ae. albopictus* populations. These associations highlight populations that are simultaneously connected through commercial exchange and exhibit relatively high genetic similarity, providing hypotheses regarding potential invasion links (Figure 7).

**Figure 7.**
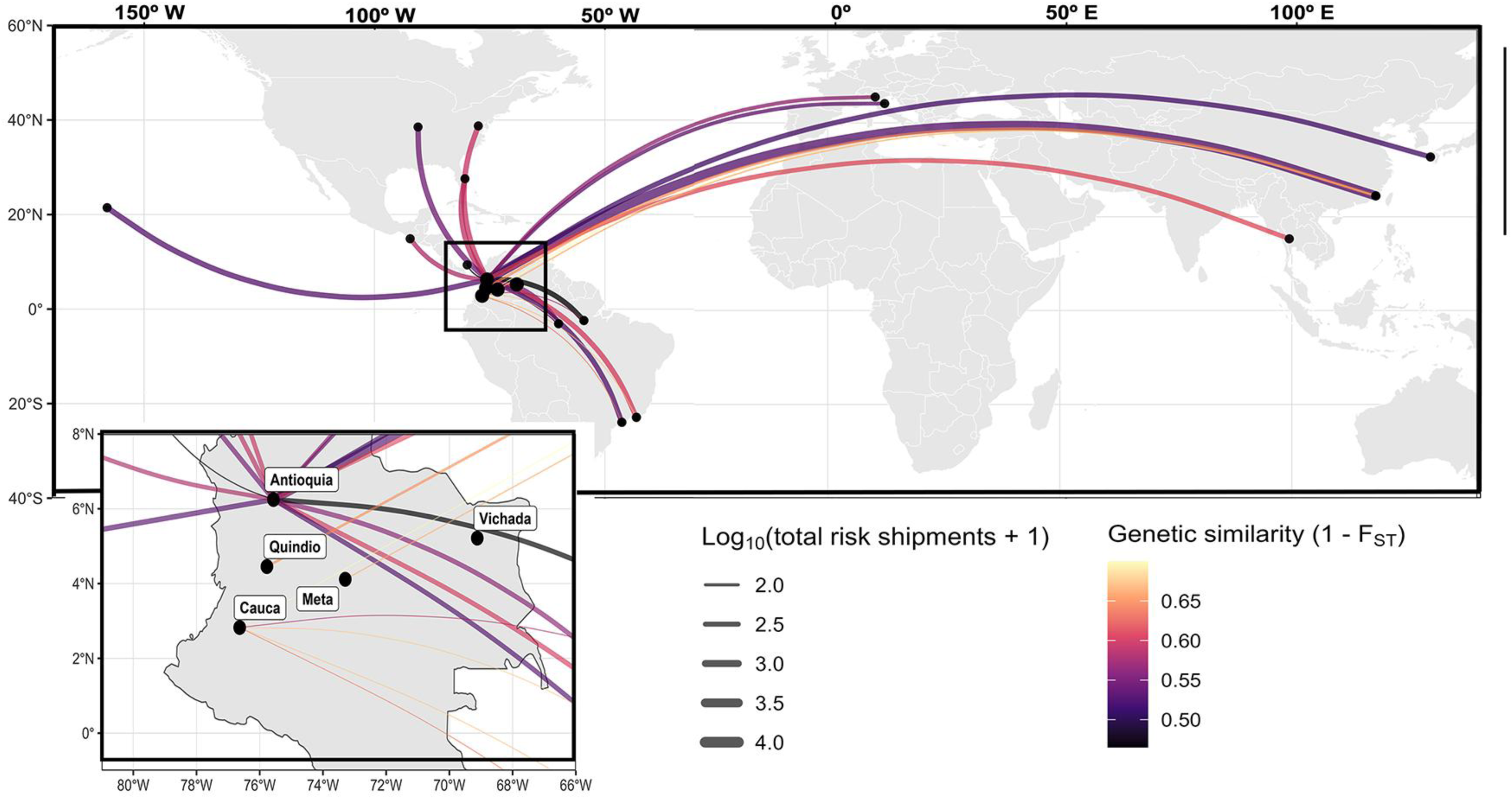
Overlap between trade connectivity and nuclear genetic similarity.

The strongest trade-genetic associations involved Asian, European, and American populations showing comparatively high genetic similarity and elevated trade connectivity with Colombian departments. Several routes converged disproportionately toward Antioquia and, to a lesser extent to Cauca, consistent with the heterogeneous trade-associated differentiation patterns identified in regression analyses. Cauca and Meta were connected through fewer high-intensity routes and exhibited lower overall microsatellite differentiation relative to several international populations. Finally, Vichada was not linked to any documented international commercial routes included in the analysis, indicating that the observed genetic affinities could not be directly related to the trade variables evaluated here.

Spatial network integrating pairwise microsatellite genetic similarity and international trade connectivity between Colombian departments and international *Ae. albopictus* populations. Route width represents shipment frequency of high-risk commodities, including used tires, new tires, and live plants, whereas route color indicates relative microsatellite genetic similarity (1 − FST) between connected populations. Black points indicate international reference populations and Colombian departments included in the analysis. The inset highlights Colombian departmental destinations and the convergence of multiple international trade-associated routes into Antioquia, Cauca, Meta, Quindío, and Vichada. Curved connections represent trade-associated connectivity links identified through the integration of shipment frequency and microsatellite genetic similarity and should not be interpreted as direct evidence of dispersal trajectories or introduction events.

Rather than supporting a single geographic affinity or a simple expansion pattern, the spatial network revealed a complex set of partially overlapping trade-genetic connections involving multiple regions and commodity categories. The coexistence of highly connected trade links with varying levels of genetic similarity highlights the heterogeneous relationships between commercial connectivity and contemporary nuclear genetic structure among Colombian departments. Overall, these analyses indicate that multiple international populations exhibit varying degrees of genetic affinity and trade connectivity with Colombian *Ae. albopictus* populations, consistent with a complex invasion scenario involving geographically diverse lineages.

### Wolbachia detection

Of the 294 *Ae.* albopictus specimens screened, *Wolbachia* infection was detected in 178 individuals (61%), although prevalence varied markedly among departments. In Antioquia, prevalence reached 100%, with high frequencies of both strains (wAlbA 95%; wAlbB 85%) and a high proportion of dual infections (88%). Cauca had the lowest prevalence (34%), with wAlbB (31%) slightly more frequent than wAlbA (24%), and fewer co-infections (23%). Meta exhibited intermediate prevalence (67%), with similar frequencies of both strains wAlbA (66%) and wAlbB (60%). In Quindío, 43% of specimens were positive, all co-infected. Vichada exhibited high prevalence (88%), with wAlbB being the dominant strain (98%) and extensive co-infection (85%). Overall, both *Wolbachia* lineages were widespread across Colombian populations, although prevalence varied markedly among departments and wAlbB generally occurred at higher frequencies than wAlbA, particularly in Vichada (Table 1).

### Effect of Wolbachia infection on host genetics

Population-level analyses, although not statistically significant, showed stronger associations between *Wolbachia* prevalence and mitochondrial than nuclear genetic metrics. Negative associations were observed between *Wolbachia* prevalence and mitochondrial differentiation (mean ΦST), particularly for wAlbB (ρ = −0.70, p = 0.23), whereas no comparable pattern was detected for nuclear differentiation (mean FST; ρ = 0.30, p = 0.68). Higher *Wolbachia* prevalence and coinfection frequencies were also associated with greater mitochondrial diversity and lower mitochondrial differentiation, with mitochondrial diversity showing strong negative correlations with mean ΦST (Hd: ρ = −0.80, p = 0.13; π: ρ = −0.90, p = 0.08). Pairwise analyses further showed stronger correlations between differences in Wolbachia coinfection prevalence and mitochondrial differentiation (ρ = 0.45, p = 0.19) or mito-nuclear discordance estimated as ΦST − FST (ρ = 0.43, p = 0.22) than with nuclear differentiation alone (ρ = −0.19, p = 0.61; Figure 8).

**Figure 8.**
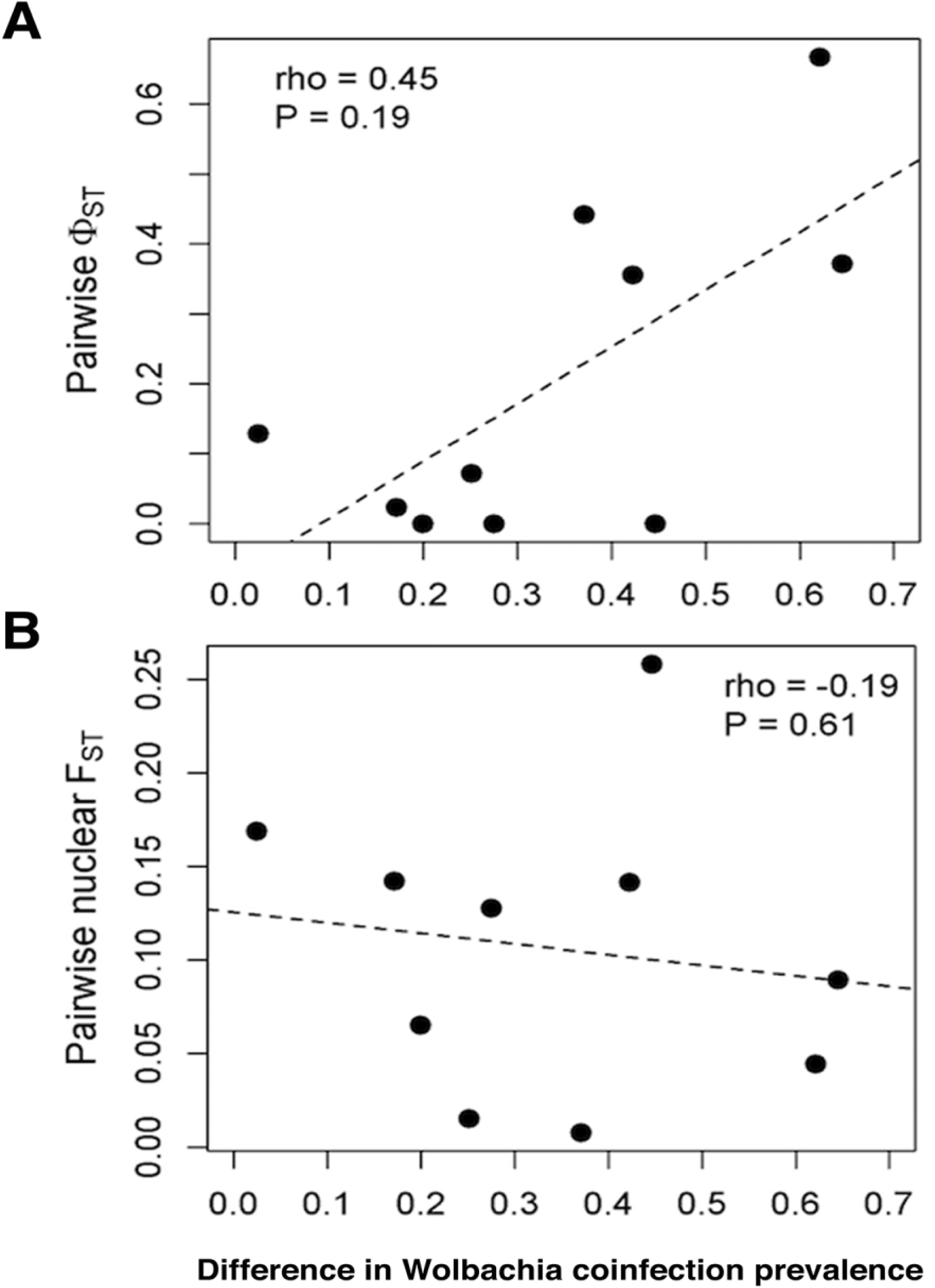
*Wolbachia* coinfection differences and mitochondrial versus nuclear population differentiation in Colombian populations of Aedes albopictus. (A) Relationship between pairwise differences in *Wolbachia* A+B coinfection prevalence and mitochondrial differentiation estimated by COI ΦST. (B) Relationship between pairwise differences in *Wolbachia* coinfection prevalence and nuclear differentiation estimated by microsatellite FST. Dashed lines represent linear regression trends. *Wolbachia* coinfection differences showed a moderate positive association with mitochondrial differentiation (ρ = 0.45, p = 0.19), whereas no comparable relationship was observed for nuclear differentiation (ρ = −0.19, p = 0.61), consistent with a predominantly maternally inherited cytoplasmic signal.

Although exploratory and limited by the small number of sampled populations, these results suggest that Wolbachia-related processes may be more closely associated with mitochondrial than nuclear patterns of population structure. The observed relationships are therefore consistent with a potential contribution of *Wolbachia* to localized mitochondrial differentiation and mito-nuclear discordance among Colombian *Ae. albopictus* populations.

### Arbovirus detection

A total of 303 larvae were tested for the presence of arboviruses, with 67 testing positive (22.1%). The positive rates were 12.2% (37) for CHIKV and 13.2% for DENV (40). Both DENV-1 and DENV-2 serotypes were detected across all surveyed departments. The highest DENV positivity was observed in Meta (23.0% of tested larvae), followed by Quindío (20.0%), Cauca (11.2%), and Vichada (11.3%) (Table 3). CHIKV was detected in three departments Cauca, Meta, and Vichada with particularly high frequencies in Meta (17.6%). No evidence of ZIKV RNA was found in any population. These findings provide molecular evidence of DENV and CHIKV circulation in *Ae. albopictus* populations across multiple Colombian departments

**Table 3.**
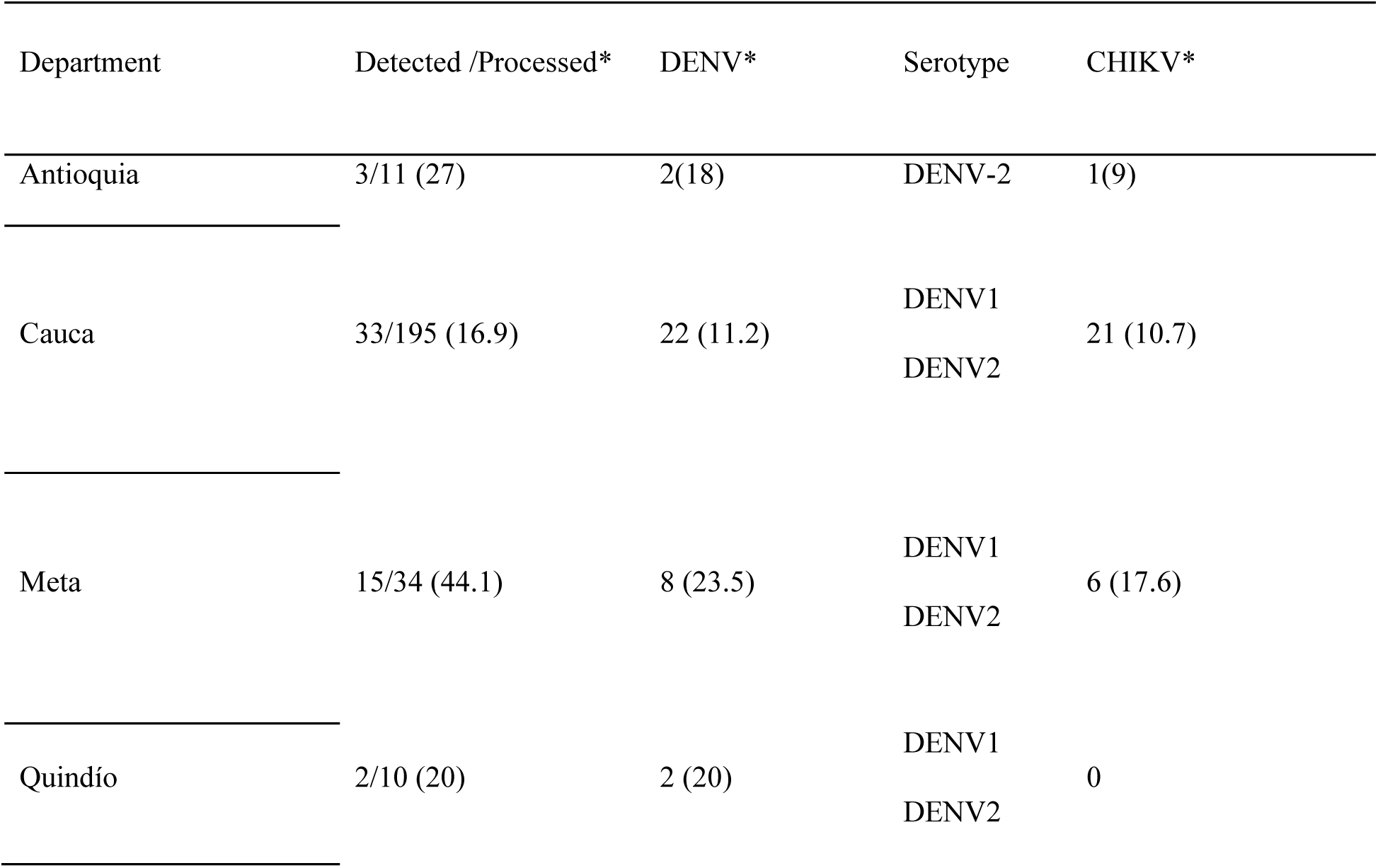
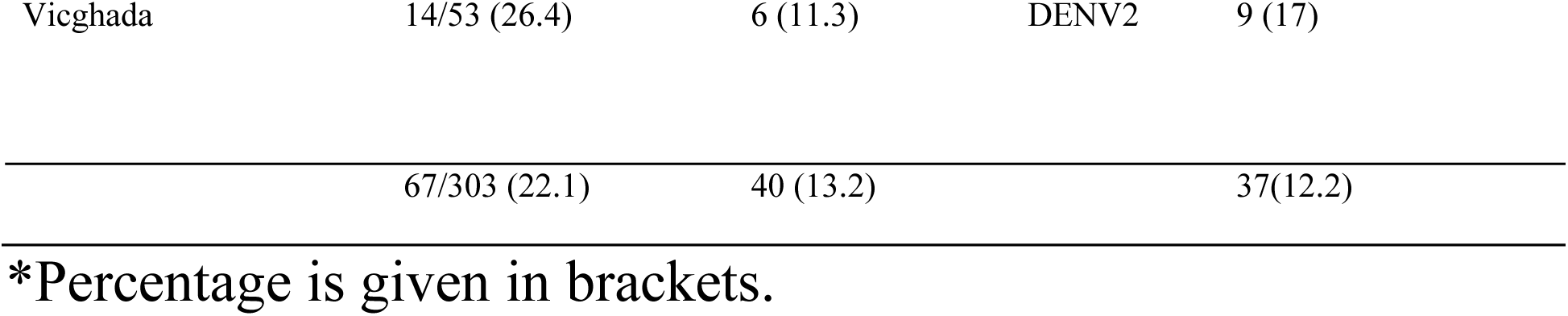
Arbovirus detection on individualized *Ae. albopictus* larvae.

| Department | Detected /Processed* | DENV* | Serotype | CHIKV* |
| --- | --- | --- | --- | --- |
| Antioquia | 3/11 (27) | 2(18) | DENV-2 | 1(9) |
| Cauca | 33/195 (16.9) | 22 (11.2) | DENV1<br>DENV2 | 21 (10.7) |
| Meta | 15/34 (44.1) | 8 (23.5) | DENV1<br>DENV2 | 6 (17.6) |
| Quindío | 2/10 (20) | 2 (20) | DENV1<br>DENV2 | 0 |
| Vicghada | 14/53 (26.4) | 6 (11.3) | DENV2 | 9 (17) |
|  | 67/303 (22.1) | 40 (13.2) |  | 37(12.2) |

\*Percentage is given in brackets.

## DISCUSSION

Our results indicate that the invasion dynamics of *Aedes albopictus* in Colombia have been shaped by multiple introductions, contemporary human-mediated connectivity, and partially discordant nuclear and mitochondrial histories. Nuclear and mitochondrial markers revealed contrasting patterns of population structure, with microsatellite differentiation more strongly associated with international trade-related connectivity and mitochondrial variation retaining stronger regional structure. The heterogeneous mito-nuclear discordance observed among departments suggests that different genomic compartments capture distinct dimensions of the invasion process. Together, these findings highlight the value of integrating population genetics, trade connectivity, and *Wolbachia* screening to understand the spread of invasive mosquito populations.

The combination of high mitochondrial haplotype diversity, low nucleotide diversity, and limited phylogenetic resolution is consistent with recent expansion following multiple introductions [10]. Although dominant haplotypes were shared among geographically distant populations, supporting recurrent connectivity, the heterogeneous distribution of mitochondrial variation among departments indicates that maternal lineages may retain regional structure despite contemporary dispersal.

Microsatellite analyses indicated that Colombian *Ae. albopictus* populations share genetic affinities with multiple international regions rather than forming a single homogeneous invasion lineage. Colombian populations showed greater nuclear affinity with populations from the USA, La Réunion, China, France, and other European localities than with several neighboring Latin American populations, including Brazil, México, and Panamá [7], a pattern similar to recurrent connectivity and repeated introductions reported in Europe and Asia [12,30]. Consistent with this, nuclear differentiation was more strongly associated with international trade intensity than mitochondrial variation, suggesting that contemporary population structure is linked to human-mediated connectivity. Although these associations cannot directly identify introduction sources or dispersal trajectories, they support a complex invasion scenario involving geographically diverse lineages. This interpretation is also consistent with early hypotheses proposing that *Ae. albopictus* expansion in Colombia was facilitated by commercial transportation corridors connecting Pacific ports with inland regions [31], while the strong trade-associated genetic signals observed in Cauca and Antioquia suggest that some Colombian populations remain embedded within broader international connectivity networks rather than representing solely the products of a simple west-to-east expansion process. More broadly, these findings suggest that invasion histories across Latin America may be more complex than often assumed, potentially involving overlapping introductions, recurrent connectivity, and regional genetic exchange among populations linked through commercial transportation networks.

Discrepancies between nuclear (FST) and mitochondrial (ΦST) differentiation revealed heterogeneous mito–nuclear discordance among Colombian *Ae. albopictus* populations, indicating that nuclear and mitochondrial markers capture partially distinct dimensions of the invasion process. Whereas microsatellite differentiation was more strongly associated with international commercial connectivity, including commodities repeatedly implicated in long-distance mosquito dispersal such as used tires and live plants [32–34], mitochondrial differentiation retained stronger associations with regional geographic structure. These contrasting patterns suggest that contemporary nuclear admixture linked to recurrent connectivity may coexist with the persistence of differentiated maternal lineages during regional expansion. Similar patterns of recurrent connectivity and gene flow have been reported in the Indian Ocean and Mediterranean regions, where commercial transportation networks maintain continuous *Ae. albopictus* movement across large geographic distances[35].

Heterogeneous *Wolbachia* prevalence and lineage composition among Colombian departments suggest complex endosymbiont dynamics that may contribute to localized mitochondrial differentiation patterns in *Ae. albopictu*s. Similar mito–nuclear decoupling patterns have been reported in Asian populations, where Wolbachia-associated mitochondrial sweeps can partially obscure underlying population structure [15]. Because both *Wolbachia* and mitochondrial DNA are maternally inherited, shifts in endosymbiont prevalence may influence mitochondrial variation through cytoplasmic incompatibility and hitchhiking effects [14,15,36,37]. Consistent with this possibility, exploratory analyses indicated stronger associations between *Wolbachia* prevalence and mitochondrial than nuclear differentiation, although these relationships were not statistically significant and should be interpreted cautiously. Together, these observations suggest that *Wolbachia*-related processes may contribute to localized mito–nuclear discordance in Colombian *Ae. albopictus* populations.

Several Colombian populations, particularly those from Cauca and Meta, showed genetic affinity with internationally distributed lineages previously associated with high CHIKV vector competence [7]. Although vector competence cannot be inferred from population genetic similarity alone, these findings highlight the potential epidemiological relevance of genetically heterogeneous *Ae. albopictus* populations established across Colombia. In addition, the detection of DENV and CHIKV RNA in larvae, together with previous reports in adults from Cauca [5], demonstrates the presence of viral markers in multiple Colombian populations and is compatible with local viral circulation or vertical maintenance mechanisms. These observations support future studies evaluating whether Colombian populations differ in arbovirus susceptibility and transmission potential. Beyond invasion dynamics, the continued expansion of *Ae. albopictus* throughout Latin America may have important epidemiological implications. Recent evidence from Brazil documenting natural yellow fever virus infection and virus isolation from *Ae. albopictus* [38] highlights growing concern that this species may contribute to arbovirus transmission at sylvatic–urban interfaces. Overall, these findings reinforce the need to consider Ae. albopictus not only as an invasive species but also as a potential component of increasingly complex arbovirus transmission systems across the region.

Several limitations should be considered when interpreting these findings. Colombian populations were analyzed primarily at the departmental level to increase analytical robustness and facilitate integration with trade datasets reported at administrative scales. Consequently, comparisons with international city-level datasets and trade-genetic associations should be interpreted as broad-scale connectivity patterns rather than direct estimates of population exchange. Mitochondrial COI provided valuable information on maternal lineage persistence and regional structure but remained constrained by single-locus resolution and potential susceptibility to Wolbachia-associated hitchhiking [15]. In contrast, microsatellite markers captured finer contemporary patterns of admixture and trade-related connectivity despite limitations related to allele binning and partial genome coverage [39,40]. The partially discordant patterns observed between markers illustrate how different genomic compartments retain complementary demographic and dispersal signals during biological invasions. Although SNP-based approaches may provide higher resolution[30], microsatellite and mitochondrial markers remain useful tools for surveillance-oriented studies of recent invasions and human-mediated dispersal.

Overall, our results indicate that the invasion dynamics of *Ae. albopictus* in Colombia have been shaped by multiple introductions, contemporary human-mediated connectivity, and partially discordant nuclear and mitochondrial histories. By integrating population genetics, *Wolbachia* screening, and trade-related connectivity analyses, we show that different genomic compartments retain complementary signatures of the invasion process. These findings highlight the value of integrative eco-genetic approaches for understanding how biological and anthropogenic processes contribute to the spread of invasive mosquito vectors.

## ACKNOWLEDGEMENTS

We want to thank the administrative staff of Secretaría de Salud del Cauca, Secretaría de Salud del Meta, Secretaría de Salud del Vichada, and Secretaría de Salud de Armenia. We want to give special thanks to our fieldworkers. To the local community from the sampled municipalities, for receiving us and allowing us to carry out our research.

## Supplementary figures

**Figure S1.**
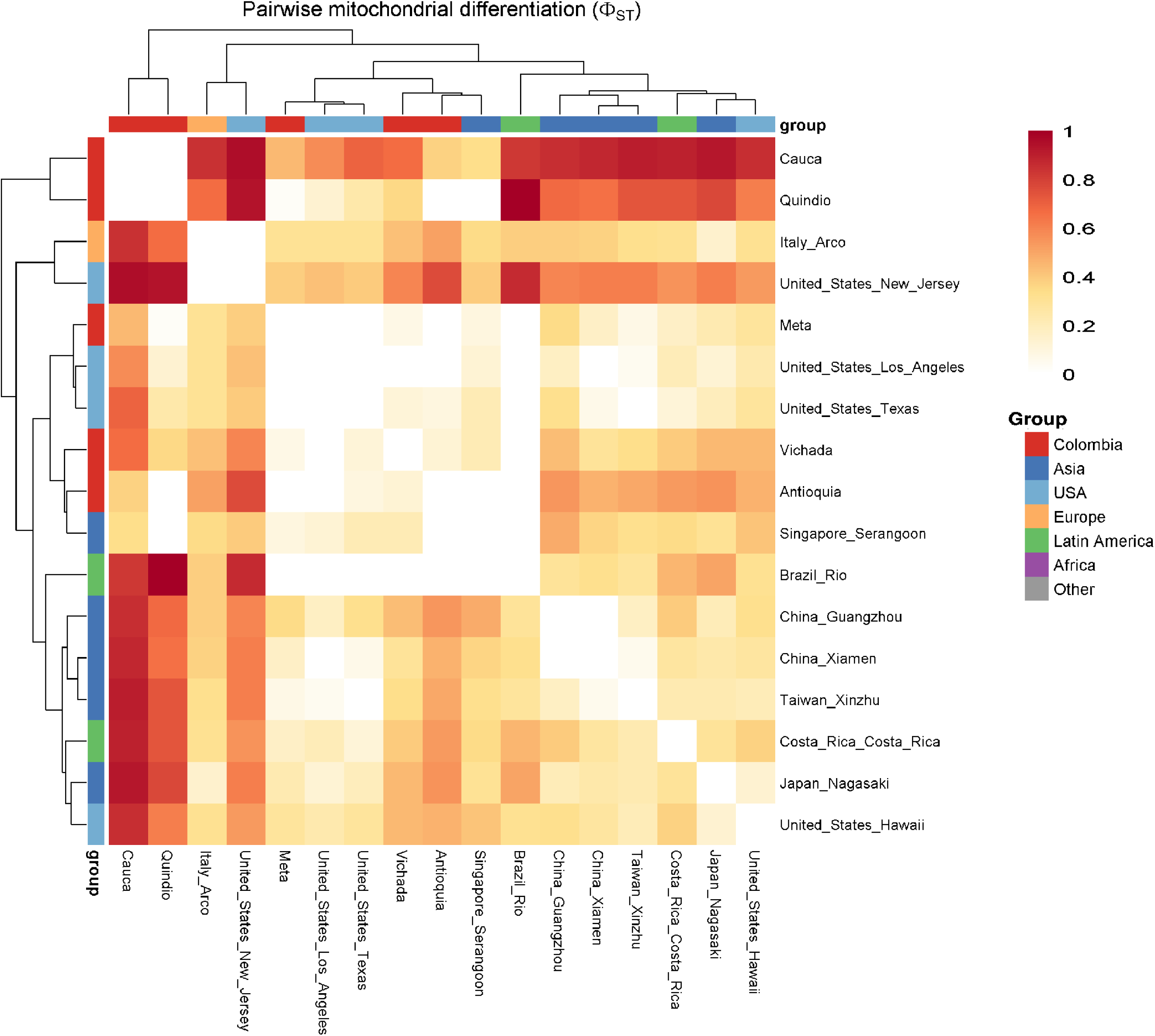
Pairwise mitochondrial differentiation among Colombian and international Aedes albopictus populations. Heatmap showing pairwise mitochondrial genetic differentiation (ΦST) estimated from COI sequences among Colombian departments and international reference populations. Hierarchical clustering was performed using pairwise ΦST distances to visualize similarities in mitochondrial lineage composition. Warmer colors indicate higher mitochondrial differentiation, whereas lighter colors indicate lower differentiation. Colored bars denote the geographic region associated with each population. The matrix illustrates heterogeneous mitochondrial relationships among Colombian populations and international lineages.

**Figure S2.**
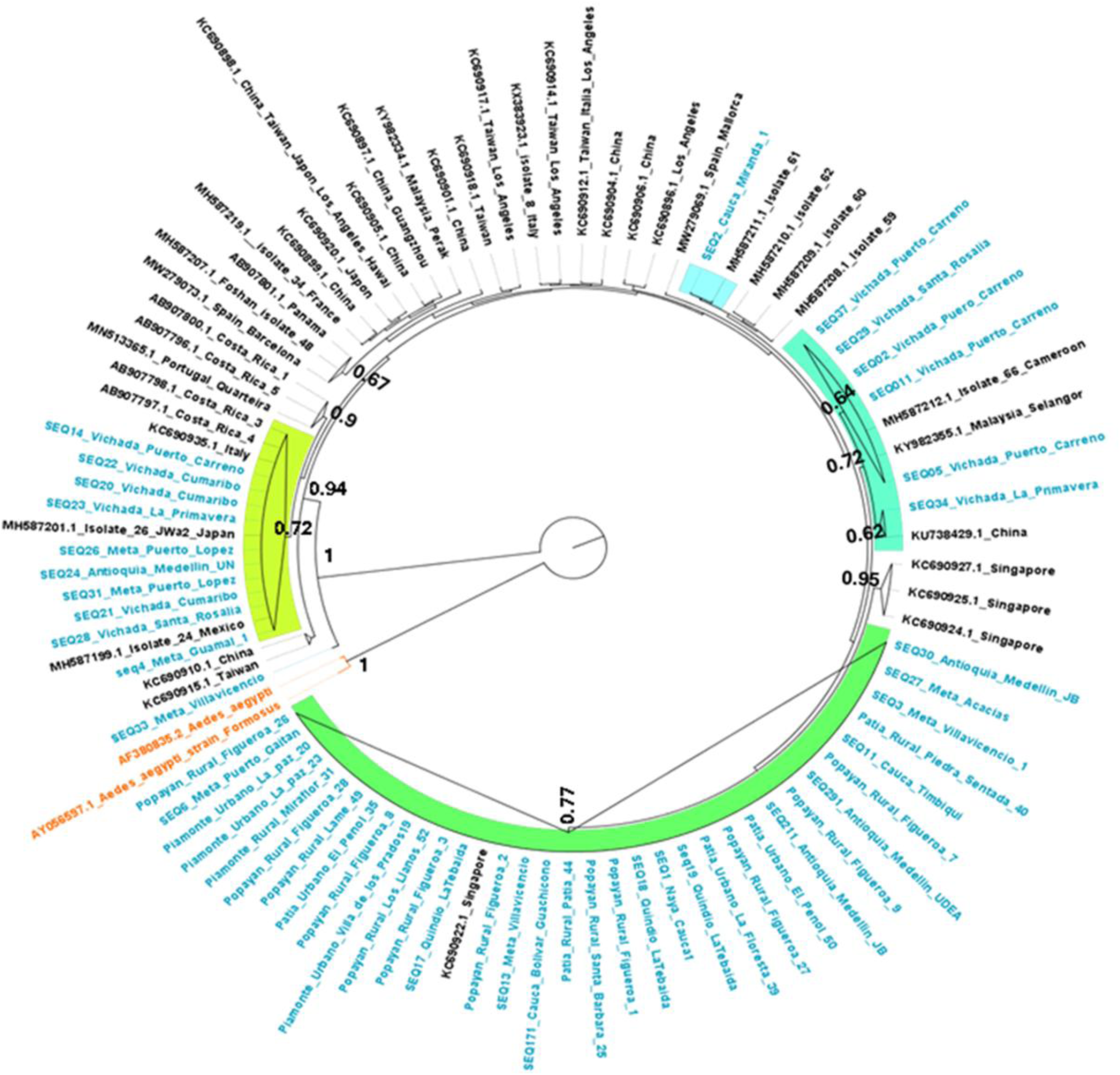
Bayesian phylogenetic reconstruction based on mitochondrial COI sequences. Bayesian phylogenetic tree inferred from a 1,350-bp mitochondrial COI alignment including Colombian and international *Aedes albopictus* sequences. Branch labels indicate posterior probability support values. Tip labels are colored according to geographic origin. Most haplotypes are separated by short branch lengths and the backbone remains poorly resolved, consistent with shallow mitochondrial divergence and recent demographic expansion.

**Figure S3.**
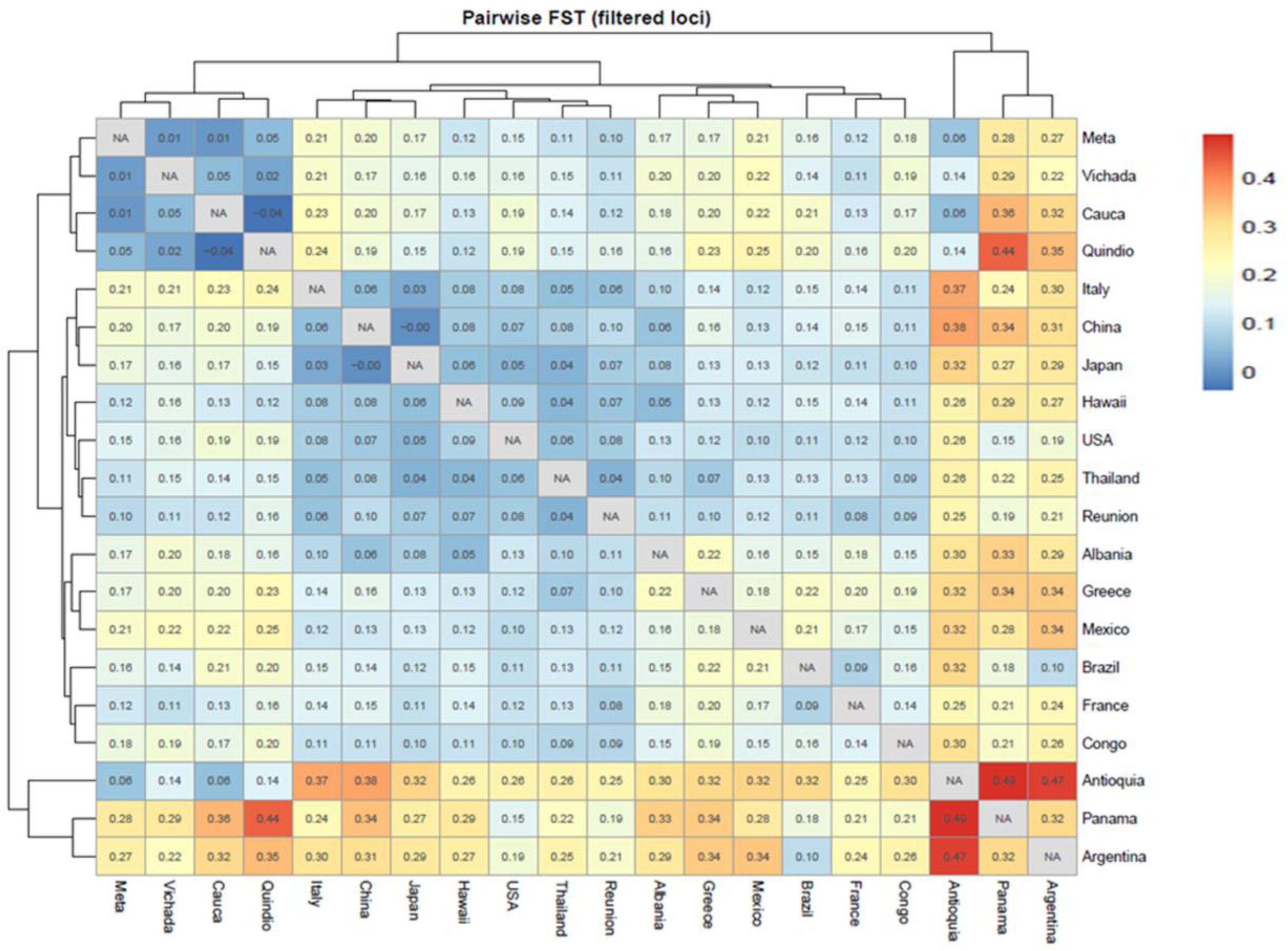
Pairwise nuclear genetic differentiation inferred from microsatellite markers. Heatmap of pairwise microsatellite differentiation (FST) calculated using the filtered eight-locus dataset after exclusion of Albmic15, Albmic6, and Albmic3. Hierarchical clustering was performed using pairwise FST distances. Warmer colors indicate greater nuclear differentiation and cooler colors indicate lower differentiation. The matrix highlights heterogeneous nuclear relationships among Colombian departments and international reference populations.

**Figure S4.**
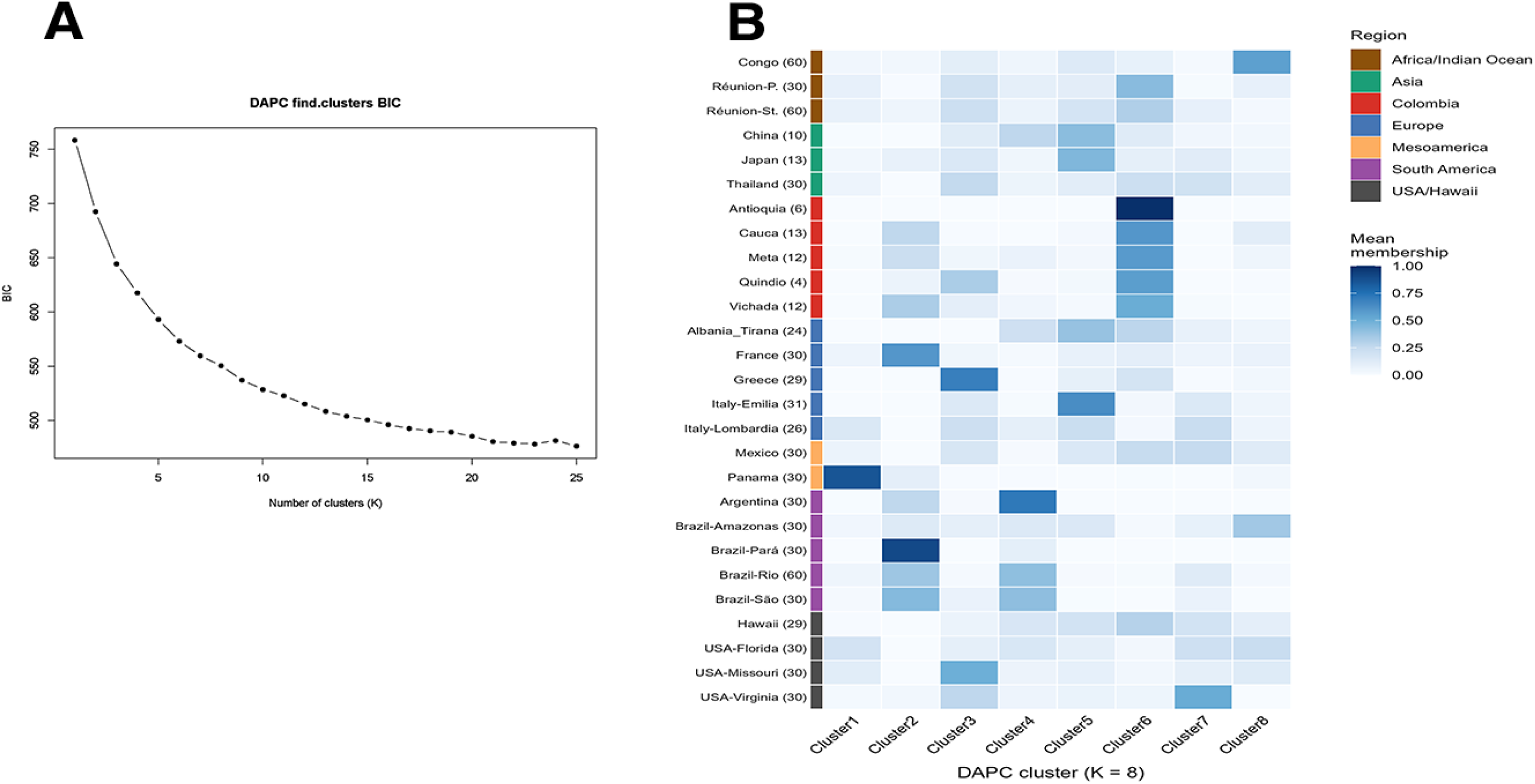
DAPC clustering and ancestry composition of international Aedes albopictus populations. (A) Bayesian Information Criterion (BIC) values obtained during DAPC clustering analyses across increasing numbers of genetic clusters (K). The selected clustering solution corresponded to K = 8. (B) Heatmap showing the proportional membership of each sampled population to the eight inferred DAPC clusters. Populations are grouped according to geographic region. Color intensity represents the relative contribution of each genetic cluster to individual populations.

**Figure S5.**
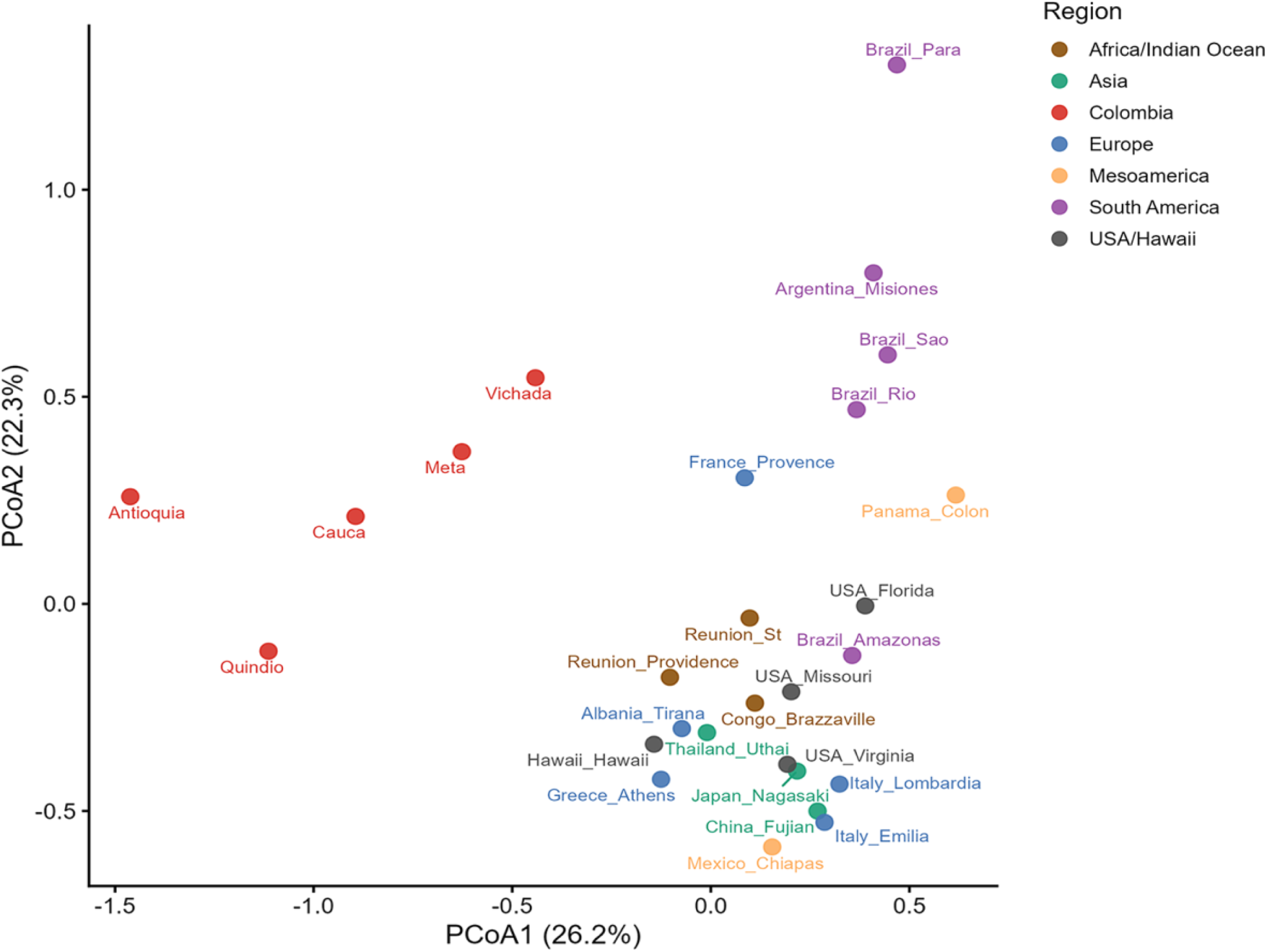
Principal Coordinate Analysis (PCoA) of microsatellite genetic variation. Principal Coordinate Analysis based on pairwise microsatellite genetic distances among Colombian and international *Aedes albopictus* populations. The first two axes explain 26.2% and 22.3% of the total genetic variation, respectively. Points are colored according to geographic region. Colombian populations occupy a partially differentiated position while maintaining proximity to populations from multiple geographic regions, consistent with mixed ancestry and complex introduction histories.

**Figure S6.**
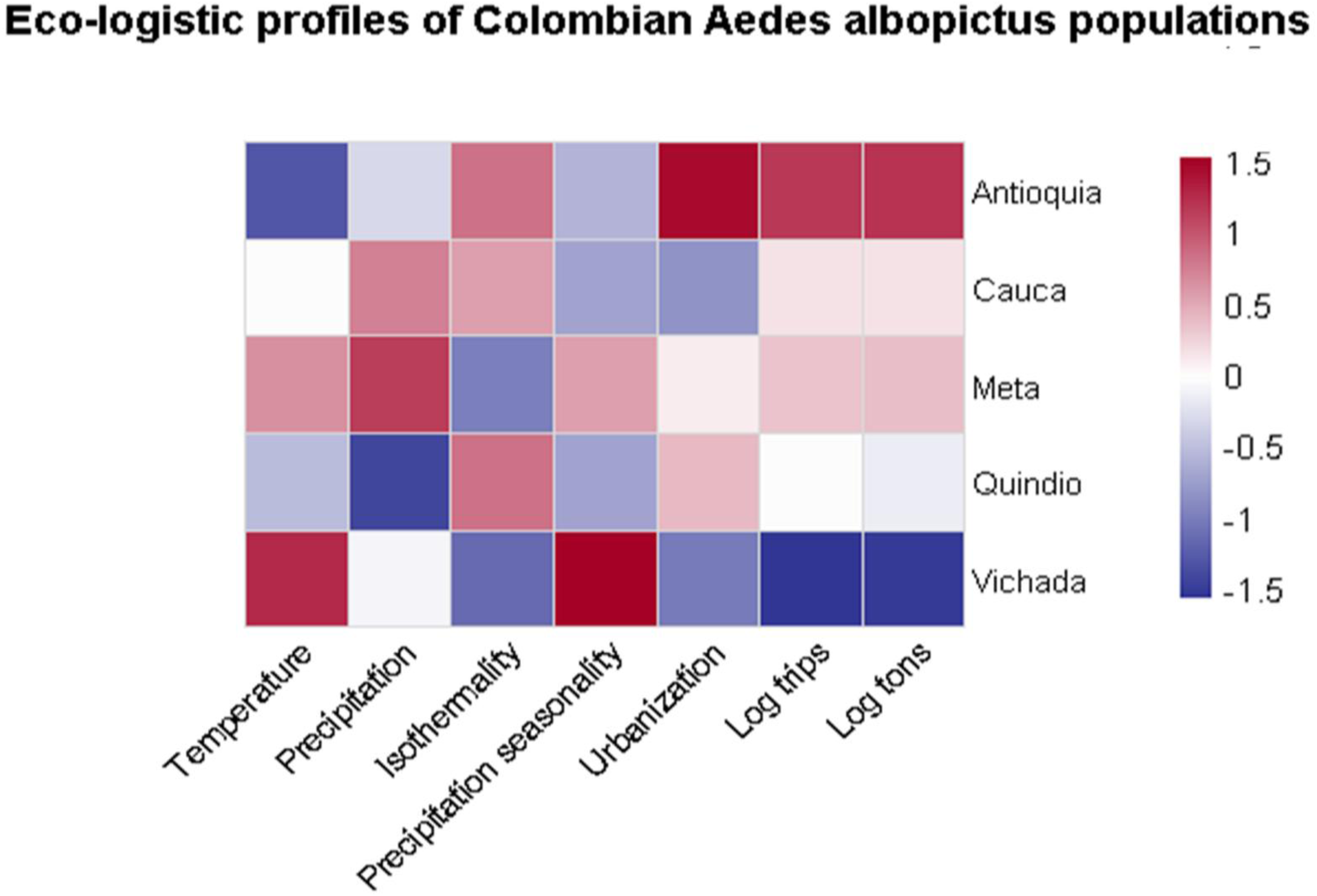
Eco-logistic profiles of Colombian Aedes albopictus populations. Heatmap summarizing standardized environmental and anthropogenic characteristics associated with sampled Colombian departments. Variables include temperature, precipitation, precipitation seasonality, urbanization, cargo trips, and cargo tonnage. Values are standardized (z-scores) to facilitate comparison among departments. Positive values indicate above-average conditions relative to the study dataset, whereas negative values indicate below-average values. The figure illustrates the environmental and logistical heterogeneity occupied by Colombian *Aedes albopictus* populations.

**Figure S7.**
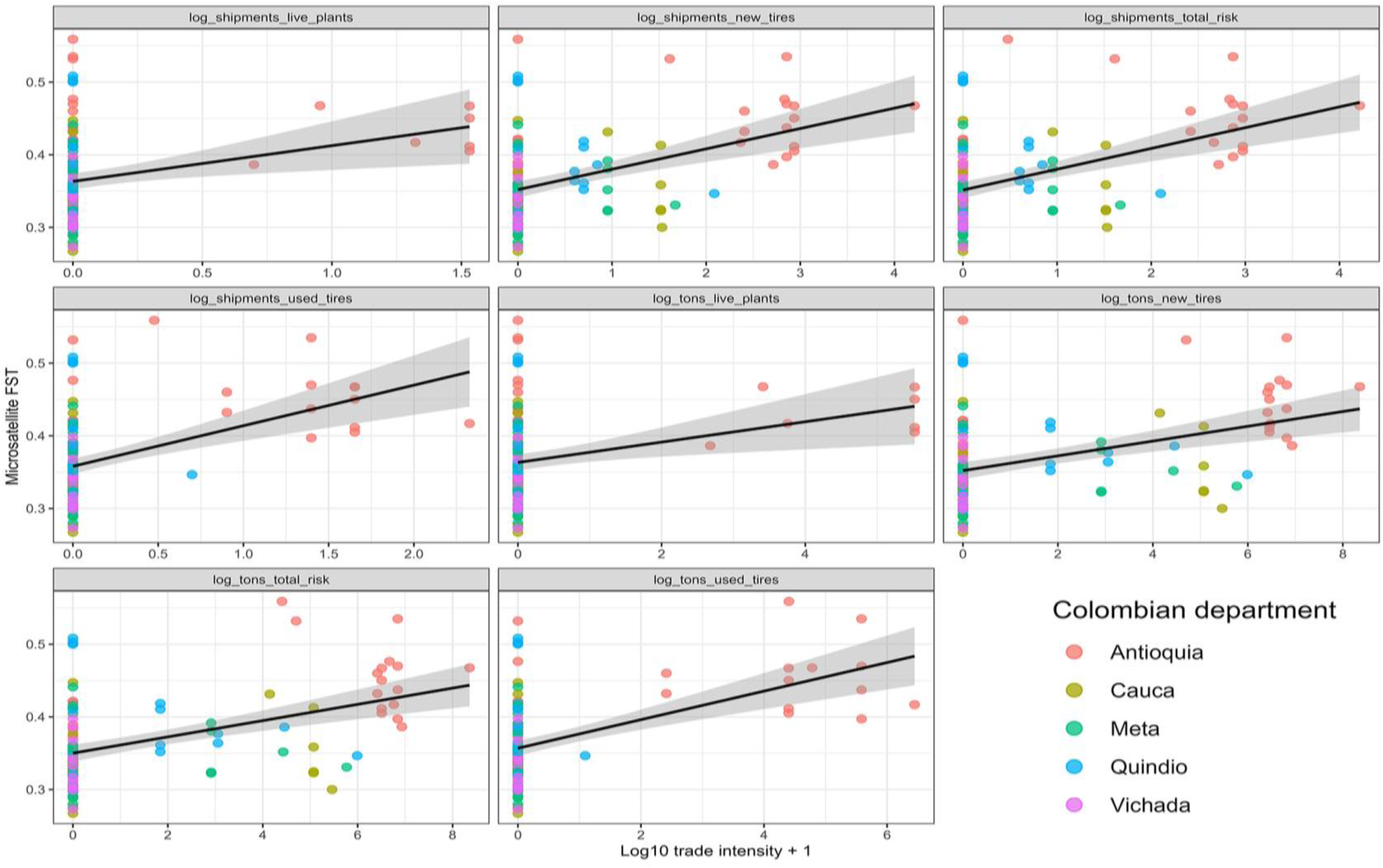
Trade-associated predictors of microsatellite genetic differentiation. Relationships between pairwise microsatellite differentiation (FST) and international trade intensity estimated using shipment frequency and import tonnage metrics for high-risk commodities, including used tires, new tires, and live plants. Each point represents an international–Colombian population comparison and colors indicate Colombian departments. Solid lines represent fitted linear models with 95% confidence intervals. Trade variables were log10-transformed prior to analysis.

**Figure S8.**
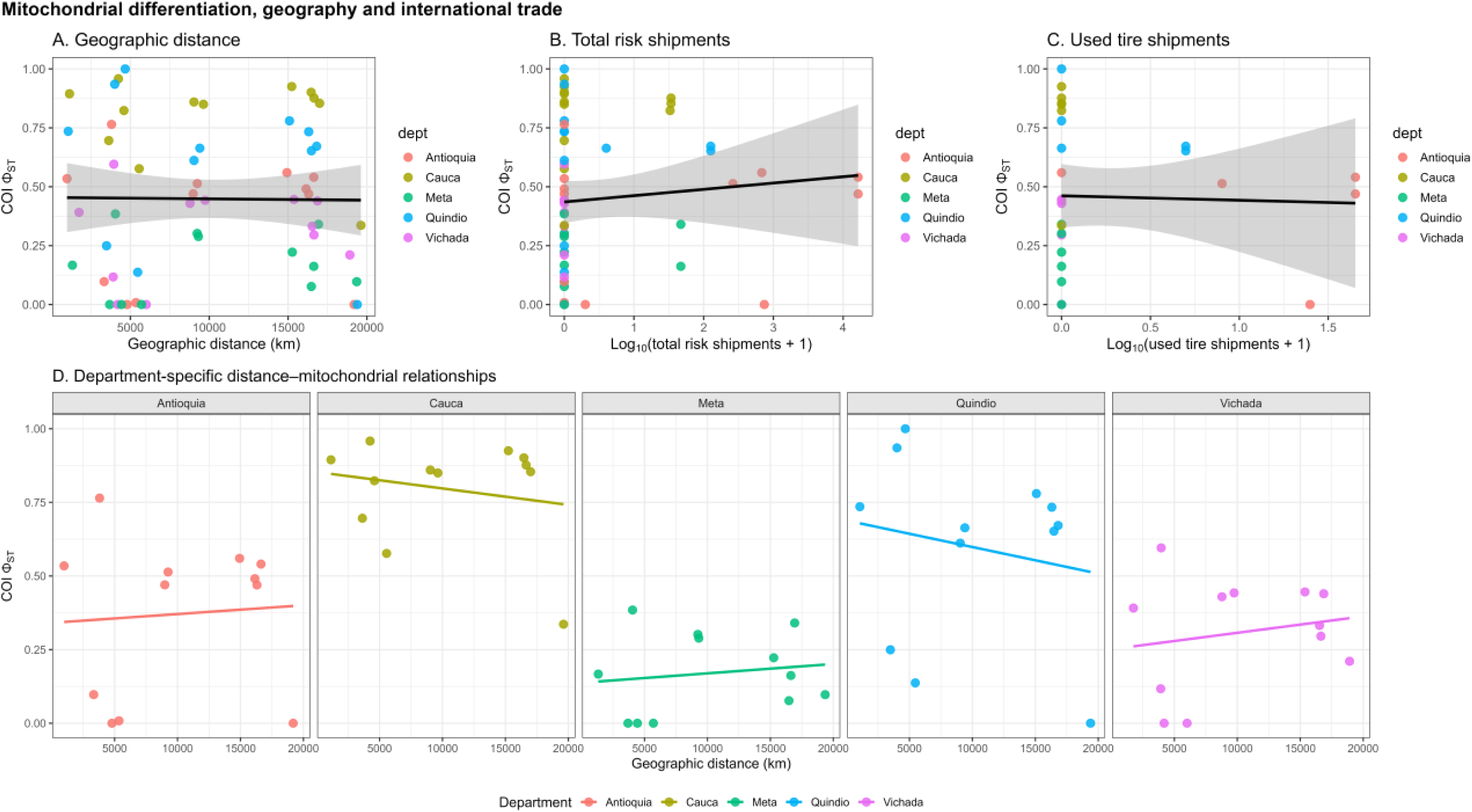
Mitochondrial differentiation in relation to geographic distance and trade-associated variables. (A) Pairwise mitochondrial differentiation (ΦST) plotted against geographic distance. (B–C) Relationships between ΦST and trade-associated shipment metrics, including total-risk shipments and used tire shipments. (D) Department-specific relationships between geographic distance and ΦST. Points are colored according to Colombian department and shaded areas represent 95% confidence intervals. These analyses illustrate the comparatively weak association between mitochondrial differentiation and trade-related variables relative to patterns observed for microsatellite markers.

## REFERENCES

1. Farooq Z, Segelmark L, Rocklöv J, Lillepold K, Sewe MO, Briet OJT, et al. Impact of climate and Aedes albopictus establishment on dengue and chikungunya outbreaks in Europe: a time-to-event analysis. Lancet Planet Heal. 2025;9: e374–e383. doi:10.1016/S2542-5196(25)00059-2

2. Bonizzoni M, Gasperi G, Chen X, James AA. The invasive mosquito species Aedes albopictus: current knowledge and future perspectives. Trends Parasitol. 2013;29: 460–468. 10.1016/j.pt.2013.07.003

3. Aguirre-Obando OA, Navarro-Silva MA. How much is known about the genetic diversity of the Asian tiger mosquito? A systematic review. Rev la Univ Ind Santander Salud. 2017;49: 422–437. doi:10.18273/revsal.v49n3-2017001

4. Mordecai EA, Cohen JM, Evans M V., Gudapati P, Johnson LR, Lippi CA, et al. Detecting the impact of temperature on transmission of Zika, dengue, and chikungunya using mechanistic models. PLoS Negl Trop Dis. 2017;11. doi:10.1371/journal.pntd.0005568

5. Mantilla-Granados JS, Montilla-López K, Sarmiento-Senior D, Chapal-Arcos E, Velandia-Romero ML, Calvo E, et al. Environmental and anthropic factors influencing Aedes aegypti and Aedes albopictus (Diptera: Culicidae), with emphasis on natural infection and dissemination: Implications for an emerging vector in Colombia. PLoS Negl Trop Dis. 2025;19: e0012605. doi:10.1371/journal.pntd.0012605

6. Ryan SJ, Carlson CJ, Mordecai EA, Johnson LR. Global expansion and redistribution of Aedes-borne virus transmission risk with climate change. PLoS Negl Trop Dis. 2019;13: e0007213. doi:10.1371/journal.pntd.0007213

7. Vega-Rúa A, Marconcini M, Madec Y, Manni M, Carraretto D, Gomulski LM, et al. Vector competence of Aedes albopictus populations for chikungunya virus is shaped by their demographic history. Commun Biol. 2020;3. doi:10.1038/s42003-020-1046-6

8. Tesla B, Demakovsky LR, Mordecai EA, Ryan SJ, Bonds MH, Ngonghala CN, et al. Temperature drives Zika virus transmission: Evidence from empirical and mathematical models. Proc R Soc B Biol Sci. 2018;285. doi:10.1098/rspb.2018.0795

9. Swan T, Russell TL, Staunton KM, Field MA, Ritchie SA, Burkot TR. A literature review of dispersal pathways of Aedes albopictus across different spatial scales: implications for vector surveillance. Parasit Vectors. 2022;15: 303. doi:10.1186/s13071-022-05413-5

10. Zhong D, Lo E, Hu R, Metzger ME, Cummings R, Bonizzoni M, et al. Genetic Analysis of Invasive Aedes albopictus Populations in Los Angeles County, California and Its Potential Public Health Impact. PLoS One. 2013;8: 1–9. doi:10.1371/journal.pone.0068586

11. Goubert C, Minard G, Vieira C, Boulesteix M. Population genetics of the Asian tiger mosquito Aedes albopictus, an invasive vector of human diseases. Heredity. Nature Publishing Group; 2016. pp. 125–134. doi:10.1038/hdy.2016.35

12. Manni M, Gomulski LM, Aketarawong N, Tait G, Scolari F, Somboon P, et al. Molecular markers for analyses of intraspecific genetic diversity in the Asian Tiger mosquito, Aedes albopictus. Parasites and Vectors. 2015;8. doi:10.1186/s13071-015-0794-5

13. Sicard M, Bonneau M, Weill M. Wolbachia prevalence, diversity, and ability to induce cytoplasmic incompatibility in mosquitoes. Current Opinion in Insect Science. Elsevier Inc.; 2019. pp. 12–20. doi:10.1016/j.cois.2019.02.005

14. Hu Y, Xi Z, Liu X, Wang J, Guo Y, Ren D, et al. Identification and molecular characterization of Wolbachia strains in natural populations of Aedes albopictus in China. Parasit Vectors. 2020;13: 28. doi:10.1186/s13071-020-3899-4

15. Ma Z, Gao J, Wang G, Zhao M, Xing D, Zhao T, et al. Effects of Wolbachia on mitochondrial DNA variation in Aedes albopictus (Diptera: Culicidae). Acta Trop. 2025;263: 107561. 10.1016/j.actatropica.2025.107561

16. Lucati F, others. Multiple introductions and {Wolbachia} infections shape {Aedes albopictus} populations in Europe. Sci Rep. 2022;12: 21874.

17. Futami K, Valderrama A, Baldi M, Minakawa N, Rodríguez RM, Chaves LF. New and Common Haplotypes Shape Genetic Diversity in Asian Tiger Mosquito Populations from Costa Rica and Panamá. J Econ Entomol. 2015;108: 761–768. doi:10.1093/jee/tou028

18. Battaglia V, Gabrieli P, Brandini S, Capodiferro MR, Javier PA, Chen X-G, et al. The Worldwide Spread of the Tiger Mosquito as Revealed by Mitogenome Haplogroup Diversity. Front Genet. 2016;7: 208. doi:10.3389/fgene.2016.00208

19. Vélez ID, Quiñones ML, Suárez M, Olano V, Murcia LM, Correa E, et al. Presencia de Aedes albopictus en Leticia, Amazonas, Colombia. Biomédica. 1998;18: 192. doi:10.7705/biomedica.v18i3.990

20. Arboleda S, Jaramillo-O N, Benavides JE, Morales AL, Matiz MI. Natural infection of {Aedes albopictus} with dengue virus in Colombia. Am J Trop Med Hyg. 2009;81: 1157–1159.

21. Pérez-Cárdenas JE, Rodr\’\iguez-Castro A, Caicedo-Quiroga L, others. Detection of arboviruses in field-collected mosquitoes in Colombia. Vector-Borne Zoonotic Dis. 2021;21: 201–210.

22. Hoyos-Polo A, Rúa-Uribe G, Wolff M. Aedes albopictus (Skuse, 1894) (Diptera: Culicidae): nuevos registros para Colombia. Revista chilena de entomología. scielocl; 2023. pp. 825–842.

23. Rueda LM. Pictorial keys for the identification of mosquitoes (Diptera: Culicidae) associated with Dengue Virus Transmission. Zootaxa. 2004;589: 1–60–1–60. doi:10.11646/ZOOTAXA.589.1.1

24. Goubert C, Minard G, Vieira C, Boulesteix M. Population genetics of the Asian tiger mosquito {Aedes albopictus}: Genetic structure, adaptive evolution, and invasive success. Mol Ecol. 2016;25: 3347–3368.

25. Goudet J. hierfstat, a package for R to compute and test hierarchical F- statistics. Mol Ecol Notes. 2005;5: 184–186.

26. Kolde R. pheatmap: Pretty Heatmaps. 2019.

27. Jombart T, Ahmed I. adegenet 1.3-1: new tools for the analysis of genome-wide SNP data. Bioinformatics. 2011;27: 3070–3071. doi:10.1093/bioinformatics/btr521

28. Garza JC, Williamson EG. Detection of reduction in population size using data from microsatellite loci. Mol Ecol. 2001;10: 305–318.

29. Calvo EP, Sánchez-Quete F, Durán S, Sandoval I, Castellanos JE. Easy and inexpensive molecular detection of dengue, chikungunya and zika viruses in febrile patients. Acta Trop. 2016;163: 32–37. doi:10.1016/j.actatropica.2016.07.021

30. Corley M, Cosme L, Armbruster P, Beebe N, Bega A, Boyer S, et al. Population Structure of the Invasive Asian Tiger Mosquito, Aedes albopictus, in Europe. Ecol Evol. 2025;15. doi:10.1002/ece3.71009

31. Cuéllar-Jiménez ME, Velásquez-Escobar OL, González-Obando R, Morales-Reichmann CA. Detección de Aedes albopictus (Skuse) (Diptera: Culicidae) en la ciudad de Cali, Valle del Cauca, Colombia. Biomedica. 2007;27: 273– 279. doi:10.7705/biomedica.v27i2.224

32. Reiter P. {Aedes albopictus} and the world trade in used tires, 1980--2000: The shape of things to come? J Am Mosq Control Assoc. 2010;26: 1–12.

33. Eritja R, Palmer JRB, Roiz D, Sanpera-Calbet I, Bartumeus F. Direct evidence of adult {Aedes albopictus} dispersal by car. Sci Rep. 2017;7: 14399.

34. Ibáñez-Justicia A, Kampen H, Werner D, Cadar D, others. Spread of {Aedes albopictus} in Europe: Lessons from monitoring and modeling approaches. Parasites \& Vectors. 2020;13: 100.

35. Delatte H, Bagny L, Brengues C, Bouetard A, Paupy C, Fontenille D. The invaders: Phylogeography of {Aedes albopictus} (Diptera: Culicidae) in the Indian Ocean with reference to Mauritius and Réunion Islands. Infect Genet Evol. 2011;11: 767–776.

36. Bourtzis K, Dobson SL, Xi Z, Rasgon JL, Calvitti M, Moreira LA, et al. Harnessing mosquito-Wolbachia symbiosis for vector and disease control. Acta Trop. 2014;132 Suppl: S150-63. doi:10.1016/j.actatropica.2013.11.004

37. Beckmann JF, Ronau JA, Hochstrasser M. A Wolbachia deubiquitylating enzyme induces cytoplasmic incompatibility. Nat Microbiol. 2017;2: 17007. doi:10.1038/nmicrobiol.2017.7

38. Bergo ES, de-Deus JT, Mucci LF, Helfstein VC, Nascimento M de JC, Rocha NRMF, et al. Yellow Fever Virus in Aedes albopictus Mosquitoes from Urban Green Area, São Paulo State, Brazil. Emerg Infect Dis. 2025;31: 2197–2199. doi:10.3201/eid3111.250692

39. Selkoe KA, Toonen RJ. Microsatellites for ecologists: A practical guide to using and evaluating microsatellite markers. Ecol Lett. 2006;9: 615–629.

40. Guichoux E, Lagache L, Wagner S, Chaumeil P, Léger P, Lepais O, et al. Current trends in microsatellite genotyping. Mol Ecol Resour. 2011;11: 591–611.

